# Identification and Interaction Analysis of Molecular Markers in Myocardial Infarction by Integrated Bioinformatics Analysis

**DOI:** 10.1101/2021.08.28.458005

**Authors:** Basavaraj Vastrad, Chanabasayya Vastrad

**Author notes:** Chanabasayya Vastrad Chanabasava Nilaya, Bharthinagar, Dharwad 580001, Karanataka, India.

## Abstract

Myocardial infarction (MI) is the leading cardiovascular diseases in worldwide, yet relatively little is known about the genes and signaling pathways involved in MI progression. The present investigation aimed to elucidate potential crucial candidate genes and pathways in MI. expression profiling by high throughput sequencing dataset (GSE132143) was downloaded from the Gene Expression Omnibus (GEO) database, which included data from 20 MI samples and 12 normal control samples. Differentially expressed genes (DEGs) were identified using t-tests in the DESeq2 R package. These DEGs were subsequently investigated by Gene Ontology (GO) and pathway enrichment analysis, a protein-protein interaction (PPI) network, modules, miRNA-hub gene regulatory network and TF-hub gene regulatory network were constructed and analyzed. Hub genes were validated by receiver operating characteristic curve (ROC) analysis. In total, 958 DEGs were identified, of which 480 were up regulated and 478 were down regulated. GO and pathway enrichment analysis results revealed that the DEGs were mainly enriched in, immune system, neuronal system, response to stimulus, and multicellular organismal process. A PPI network, modules, miRNA-hub gene regulatory network and TF-hub gene regulatory network was constructed by using Cytoscape software, and CFTR, CDK1, RPS13, RPS15A, RPS27, NOTCH1, MRPL12, NOS2, CCDC85B and ATN1 were identified as the hub genes. Our results highlight the important roles of the genes including CFTR, CDK1, RPS13, RPS15A, RPS27, NOTCH1, MRPL12, NOS2, CCDC85B and ATN1 in MI pathogenesis or therapeutic management.

## Introduction

Myocardial infarction (MI) is one of the most prevalent cardiovascular diseases in the world [1]. MI is broadly characterized as a sudden blockage of blood flow to heart and permanent damage to the heart muscle [2]. Myocardial infarction (MI) is a main cause of morbidity and mortality all over the world [3]. The potential highly risk factors for MI include genetic risk [4], older age [5], smoking [6], hypertension [7], diabetes mellitus [8] and obesity [9]. Although some prognostic biomarkers have been exploited, the overall survival of MI remains weak due to its difficulty in early detection [10–11]. It is therefore urgent to identify novel diagnostic and prognostic biomarkers, and therapeutic target for MI.

Molecular biology investigation have identified numerous biomarkers and signaling pathway that contribute to MI, including HMGA1 [12], STXBP2 [13], CCL19 and CCL21 [14], NLRP3 [15], CD14 [16], Nrf2 signaling pathway [17], Wnt signaling pathways [18], β2AR, cAMP/PKA, and BDNF/TrkB signaling pathways [19], PTEN/PI3K/Akt signaling pathway [20] and TGF-β1 signaling pathway [21]. A further study into the molecular events linked with MI is required.

One of the recent technology RNA sequencing, which allows the study of gene expression in a high throughput manner with high sensitivity, specificity and repeatability. A significant amount of data has been produced via the use of RNA sequencing and such data has been uploaded and stored in public databases. Furthermore, many bioinformatics studies on MI have been produced in recent years [22], which proved that the integrated bioinformatics techniques could help us to further investigation and better exploring the molecular mechanisms.

In this study, we identified DEGs in expression profiling by high throughput sequencing dataset GSE132143 [23] from the Gene Expression Omnibus (GEO) database (http://www.ncbi.nlm.nih.gov/geo/) [24]. We performed Gene Ontology (GO) and REACTOME functional and pathway enrichment analysis, and constructed and analyzed a protein–protein interaction (PPI) network, modules, miRNA-hub gene regulatory network and TF-hub gene regulatory network. Diagnostic values of hub genes were determined by receiver operating characteristic curve (ROC) analysis. Collectively, the findings of the current investigation highlighted hub genes and pathways that might contribute to the pathology of MI. These may provide a basis for the advancement of future diagnostic and therapeutic target for MI.

## Materials and Methods

### Data resources

GSE132143 [23] expression profiling by high throughput sequencing dataset was downloaded from the GEO database and based on GPL18573 Illumina NextSeq 500 (Homo sapiens). The datasets contained 32 samples, including 20 MI samples and 12 normal control samples.

### Identification of DEGs

DEGs between MI and normal control specimen were identified via DESeq2 package in R bioconductor [25] with |logFC|□ > 1.37 for up regulated genes and |logFC|□ < -1.89 for down regulated genes and adjust P value□ < 0.05, and Benjamini and Hochberg false discovery rate method [26]. The heatmap of DEGs in every sample of the MI and normal control group was constructed by using gplots package in R software. The volcano plot was generated by ggplot2 package in R software.

### GO and REACTOME pathway enrichment analysis of DEGs

Gene ontology (GO) (http://geneontology.org/) [27] is a framework for the model of biology, which describes gene functions and classification includes, biological process (BP), cellular component (CC) and molecular function (MF). REACTOME pathway (https://reactome.org/) [28] is a collection of databases for recognition high-level gene functions and functions of the biological system. The g:Profiler (http://biit.cs.ut.ee/gprofiler/) [29] is an online tool that provides a comprehensive set of functional annotation tools to examine the biological content behind a number of genes. In order to analyze DEGs at the functional level, GO term enrichment analysis and REACTOME pathway enrichment analysis were conducted using g:Profiler. P<0.05 was considered to indicate a statistically significant difference.

### Construction of the PPI network and module analysis

Human Integrated Protein-Protein Interaction rEference (HiPPIE) interactome (http://cbdm-01.zdv.uni-mainz.de/~mschaefer/hippie/) [30] is an online software of interactions of genes and proteins. Cytoscape 3.8.2 (http://www.cytoscape.org/) [31] was is an open-source tool for network visualization of genes and proteins. The hub genes were identified by using the plug-in Network Analyzer of the Cytoscape software, including node degree [32], betweenness centrality [33], stress centrality [34] and closeness centrality [35]. In order to find modules of the whole network, the PEWCC1 (http://apps.cytoscape.org/apps/PEWCC1) [36] plug-in of the Cytoscape software was applied.

### MiRNA-hub gene regulatory network construction

To explore the interaction among the hub genes and miRNAs, we used the miRNet database (https://www.mirnet.ca/ [37]. Furthermore, all the known and predicted miRNA-hub gene pairs were downloaded from miRNet database and the miRNA-hub gene pairs were screened out. The miRNA-hub gene regulatory network was established via Cytoscape version 3.8.2 [31].

### TF-hub gene regulatory network construction

To explore the interaction among the hub genes and TFs, we used the NetworkAnalyst database (https://www.networkanalyst.ca/) [38]. Furthermore, all the known and predicted TF-hub gene pairs were downloaded from NetworkAnalyst database and the TF-hub gene pairs were screened out. The TF-hub gene regulatory network was established via Cytoscape version 3.8.2 [31].

### Validation of hub genes by receiver operating characteristic curve (ROC) analysis

pROC package in R statistical software [39] was used to perform ROC curve analysis and to resolve the specificity and sensitivity for all the possible thresholds of the ROC curve. The area under the curve (AUC) value of the hub genes was predicted based on the ROC curve analysis.

## Results

### Identification of DEGs

By using DESeq2 package to analyze the differential expression of the expression profiling by high throughput sequencing dataset, we obtained 958 DEGs composed of 480 up regulated genes and 478 down regulated genes (Table 1). All DEGs were displayed in volcano maps (Fig. 1). Heatmap analysis showed that these genes presented differential expression profiles between MI and normal control (Fig. 2).

**Fig. 1.**
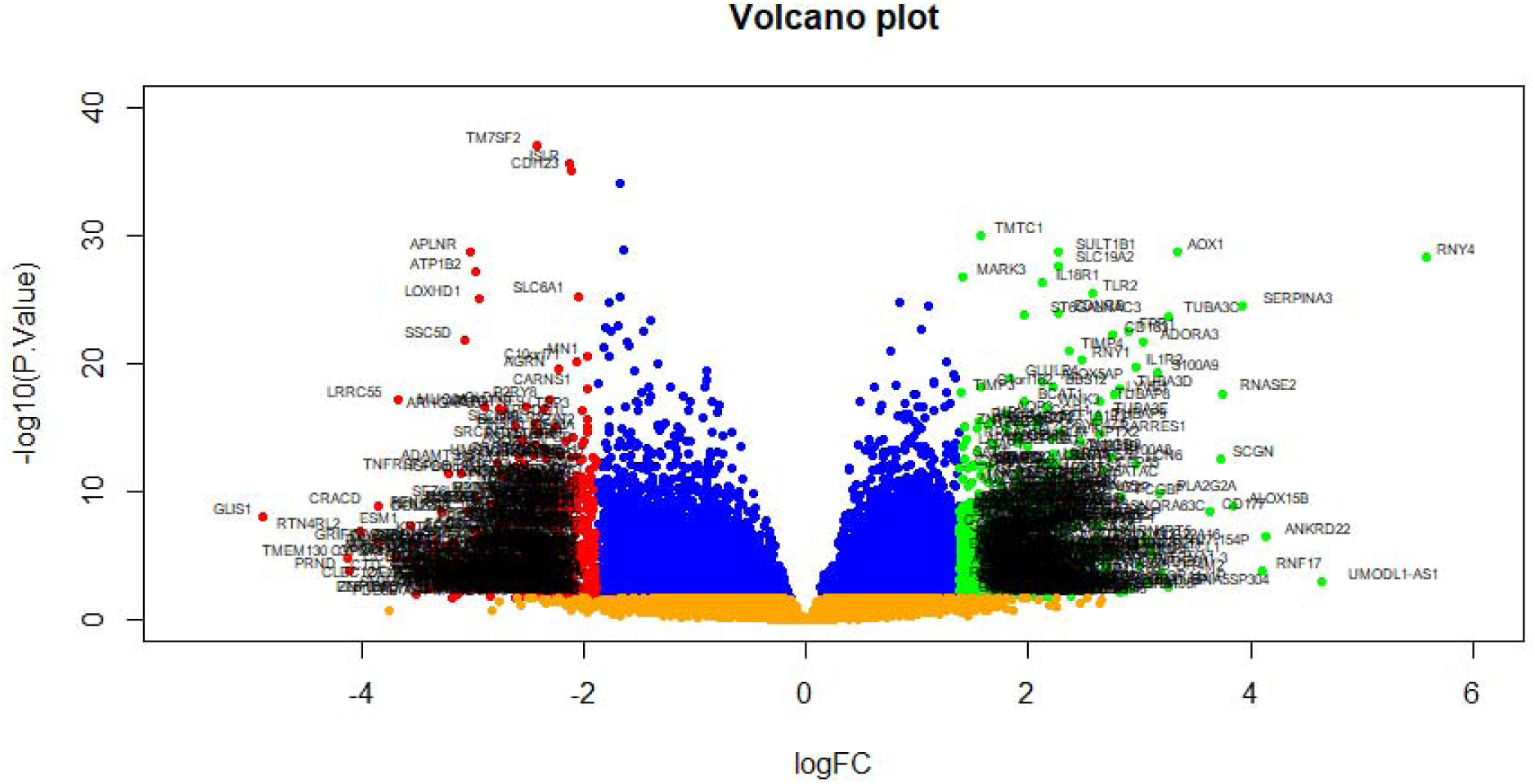
Volcano plot of differentially expressed genes. Genes with a significant change of more than two-fold were selected. Green dot represented up regulated significant genes and red dot represented down regulated significant genes.

**Fig. 2.**
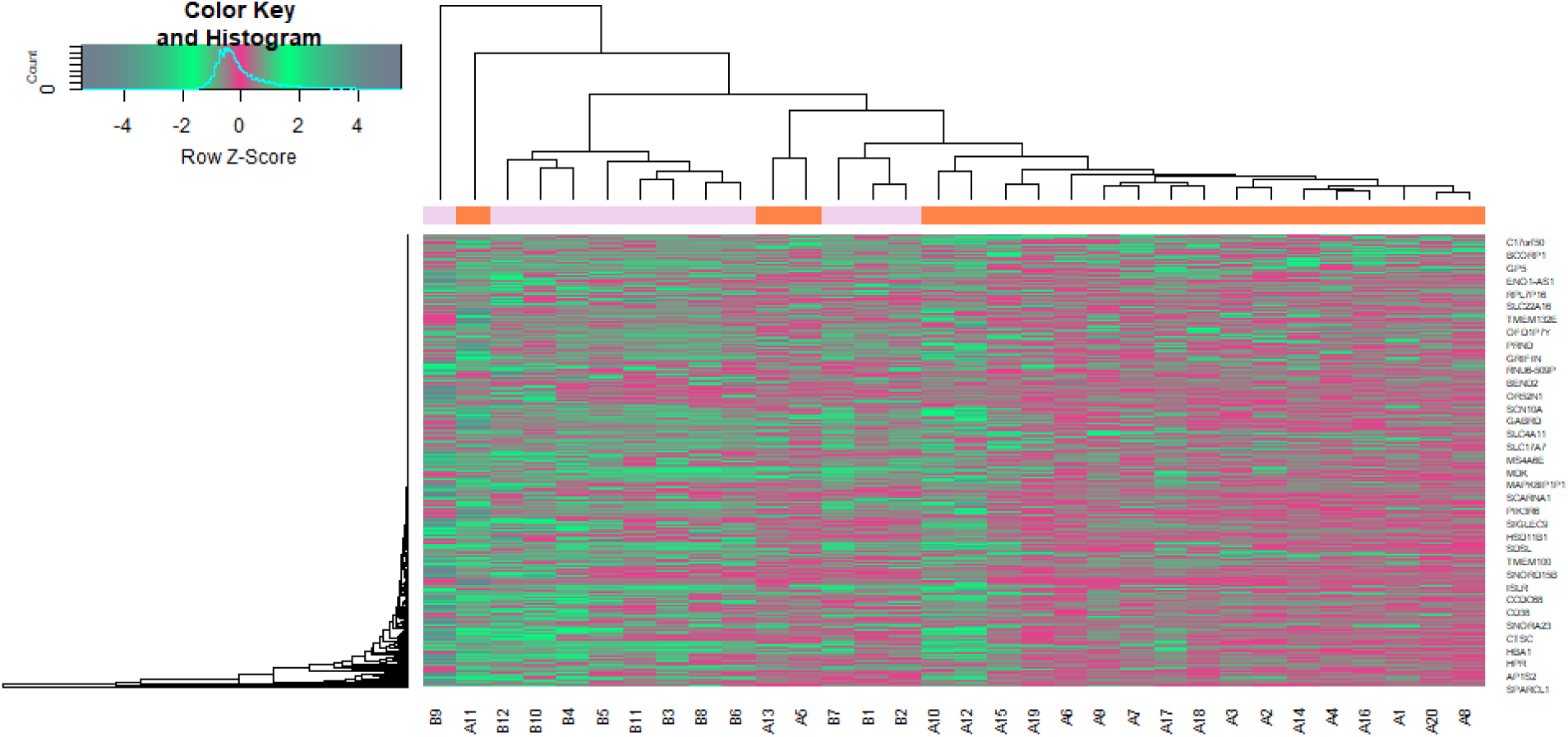
Heat map of differentially expressed genes. Legend on the top left indicate log fold change of genes. (A1 – A20= MI samples; B1 – B12 = normal control samples)

**Table 1.**
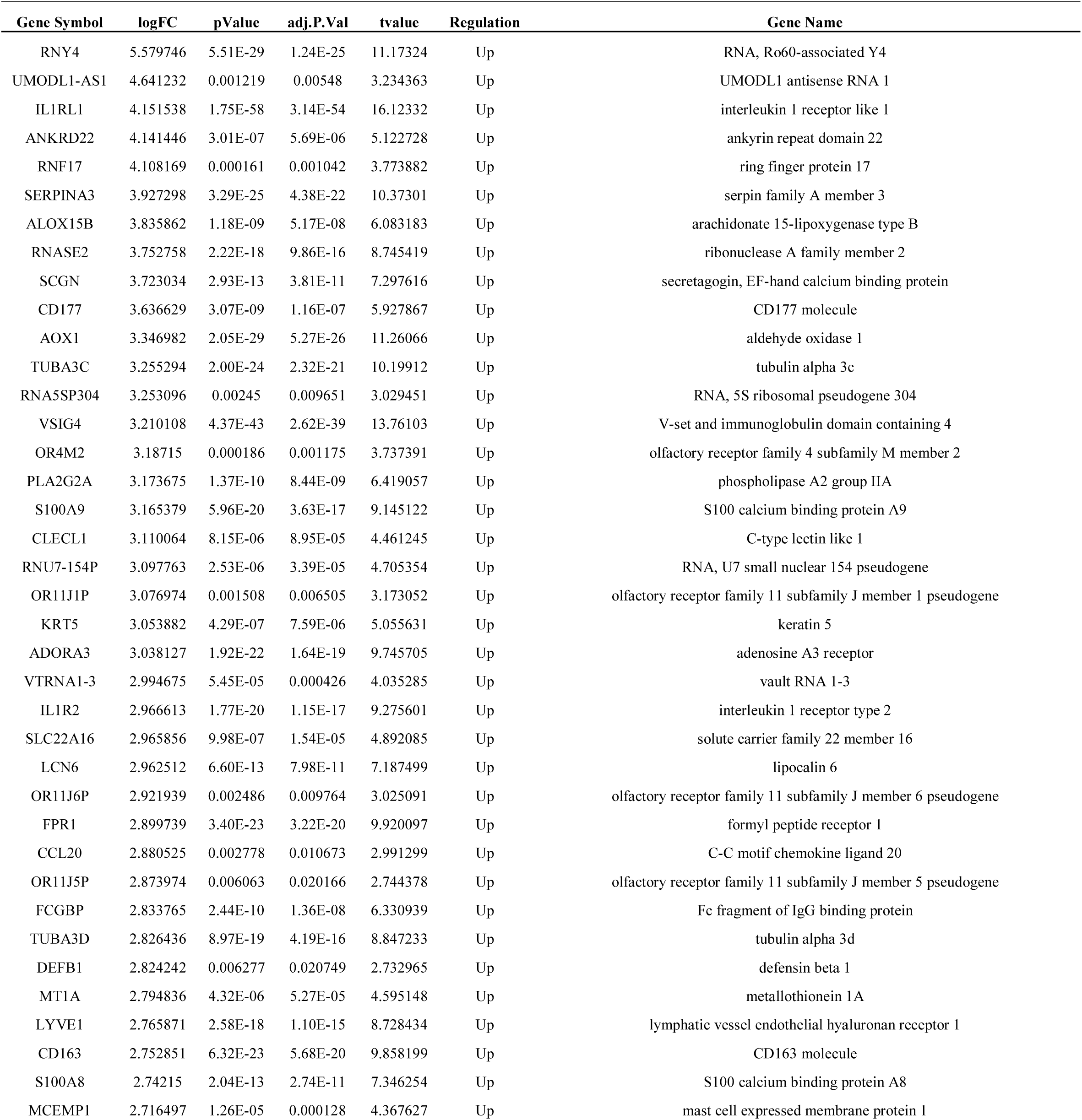
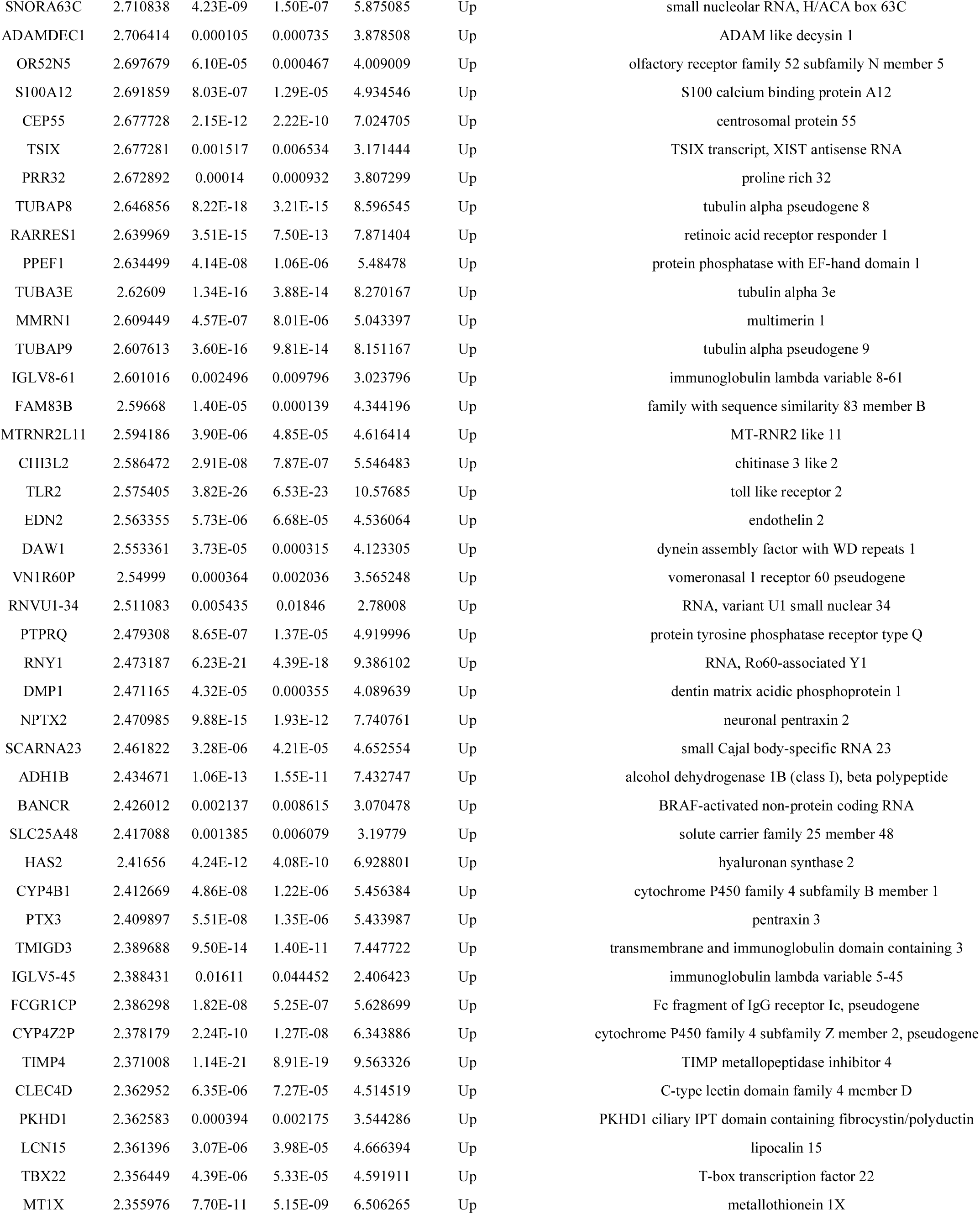
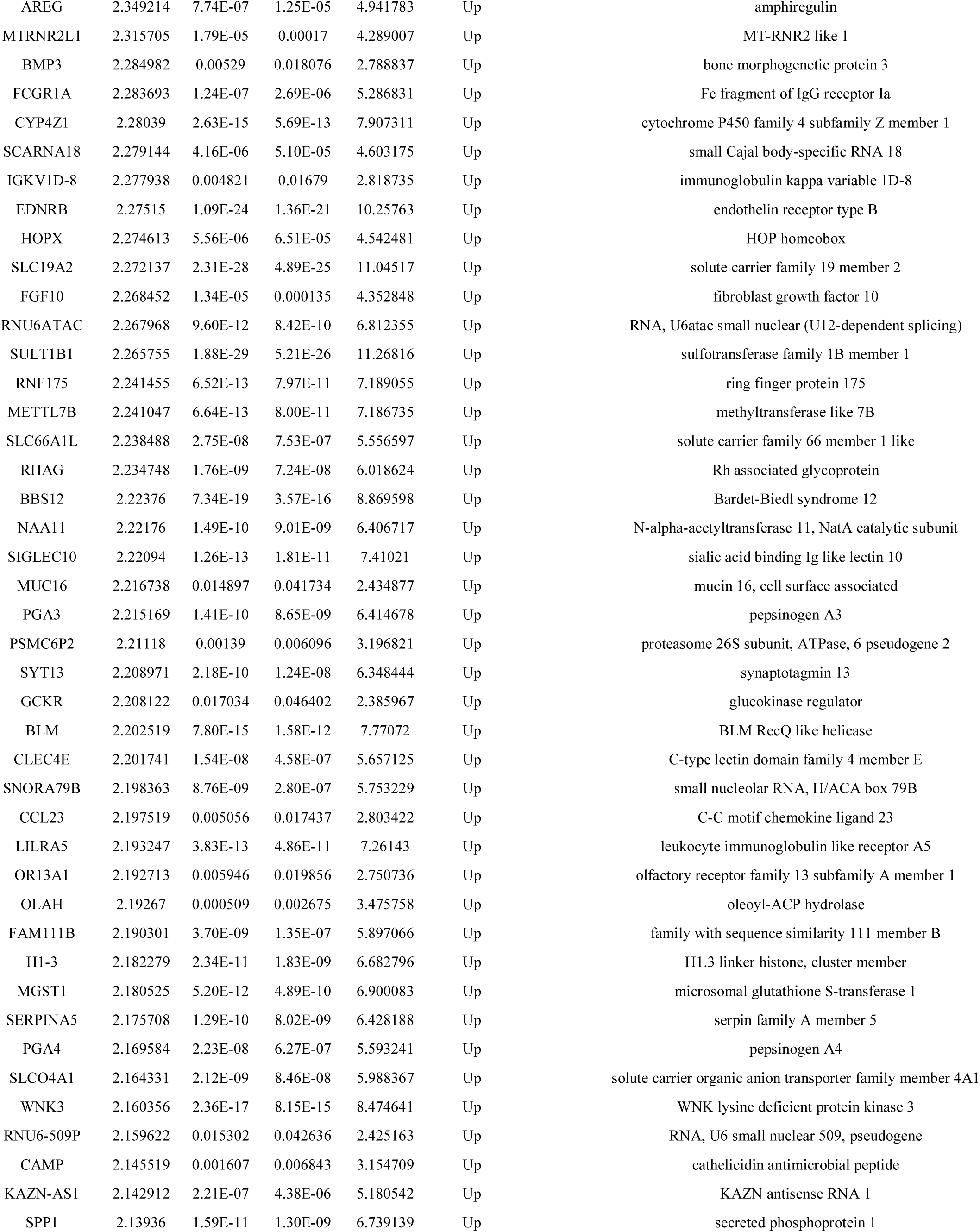
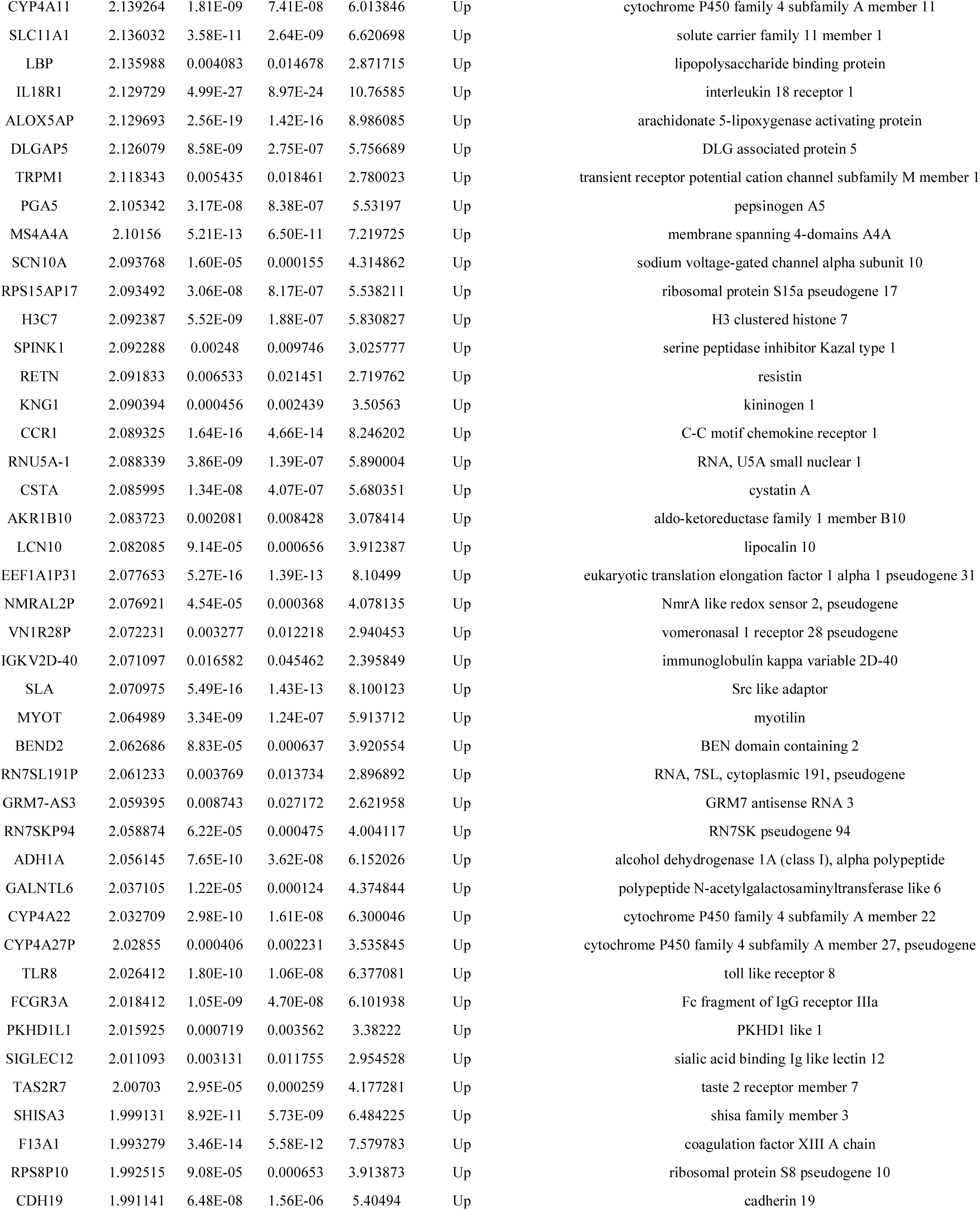
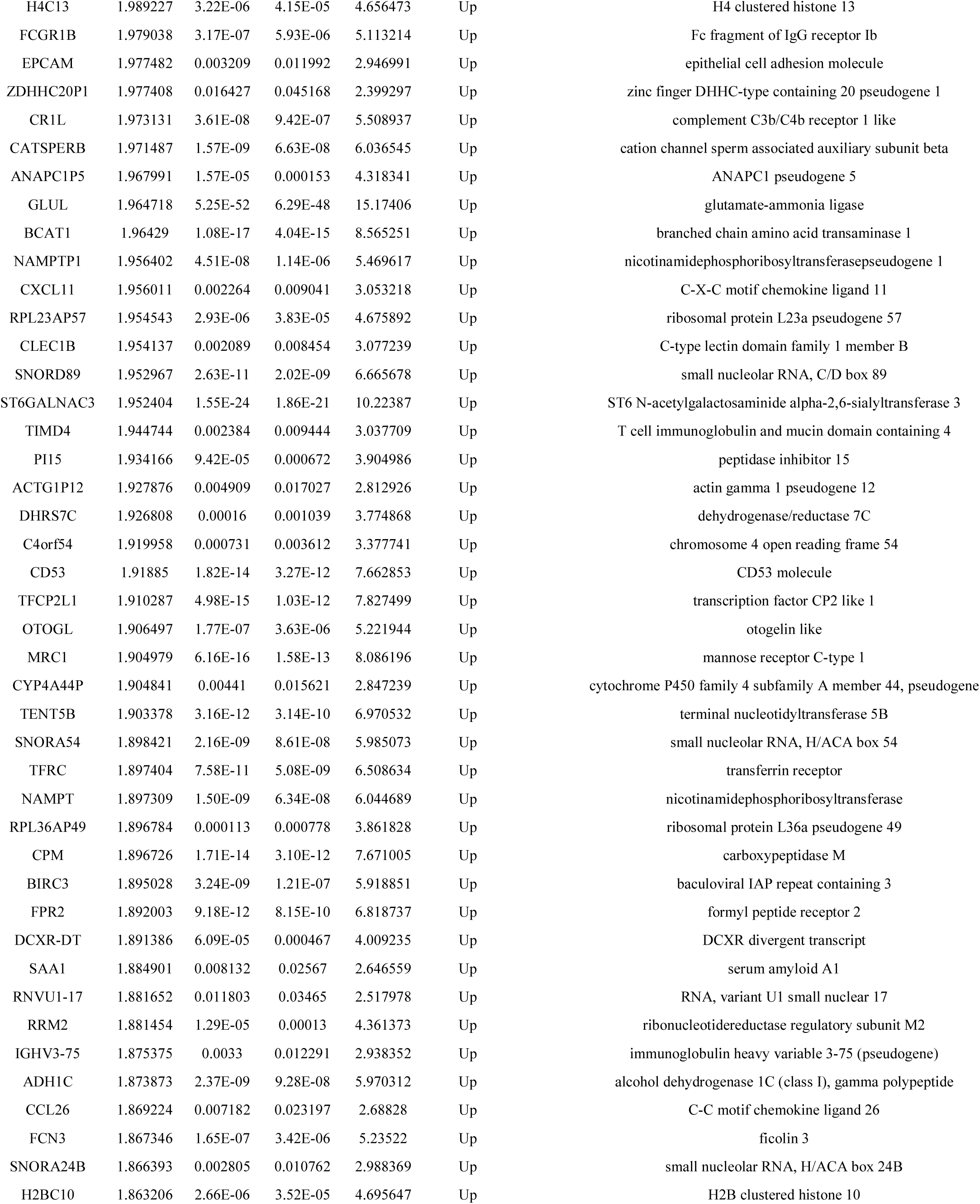
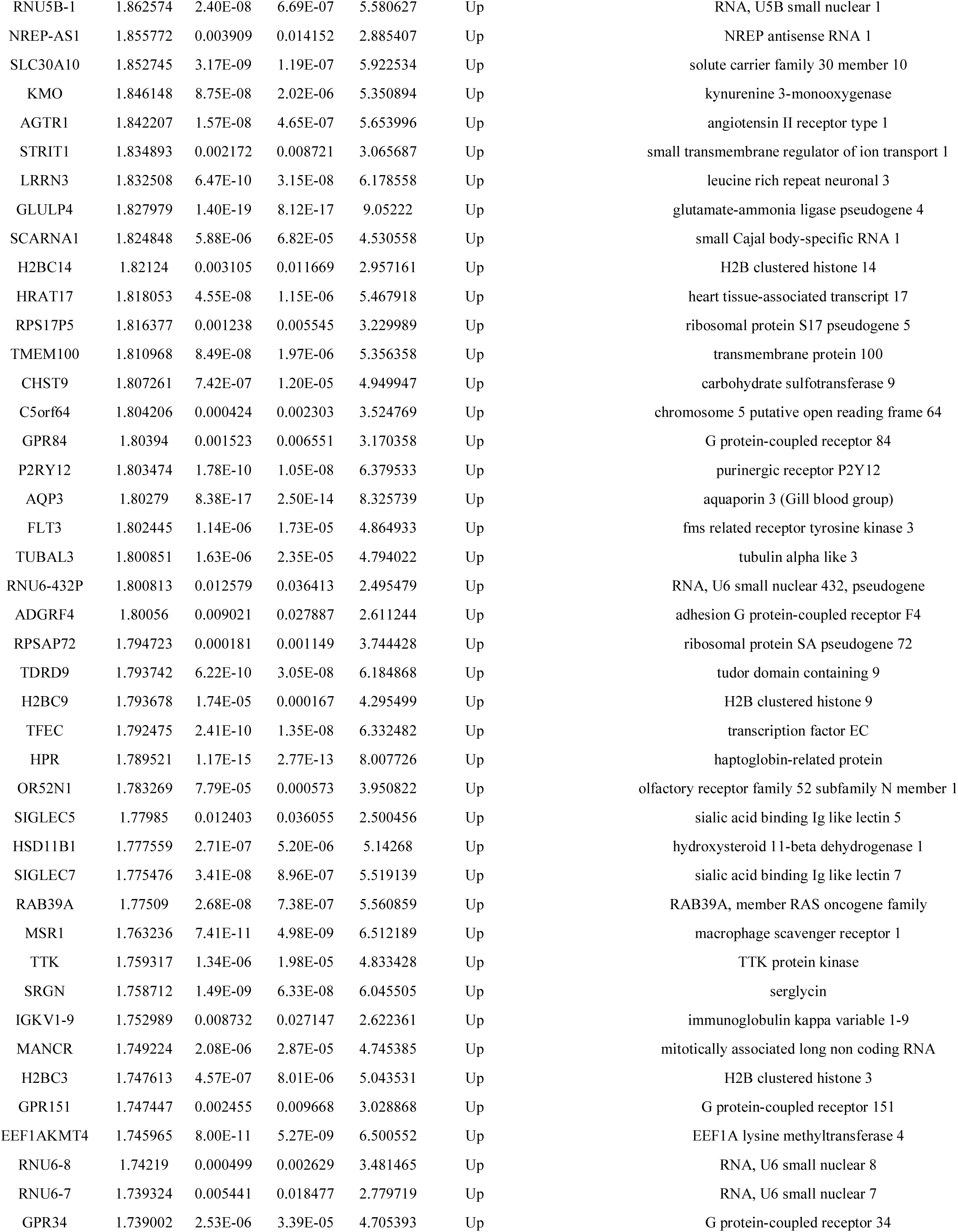
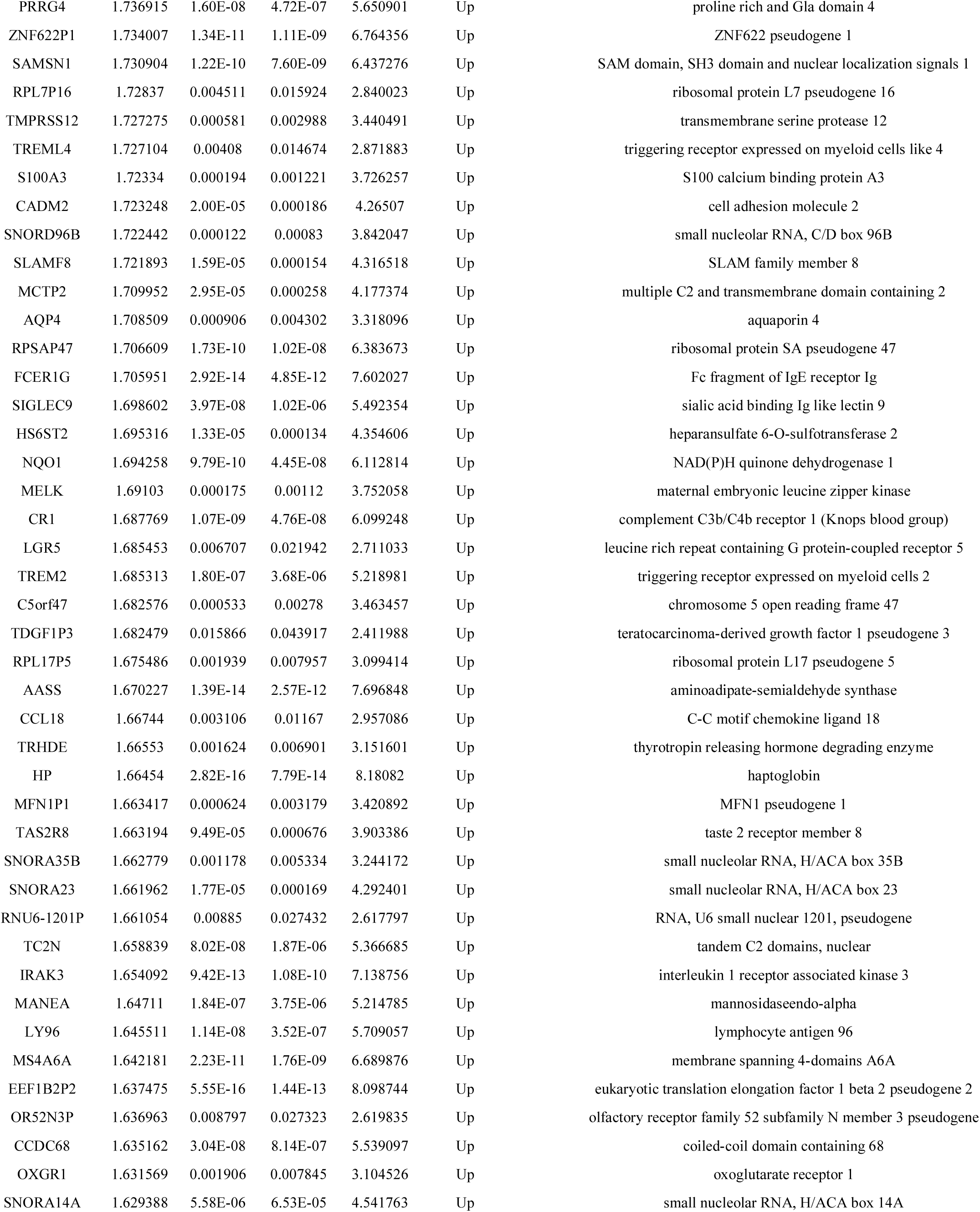
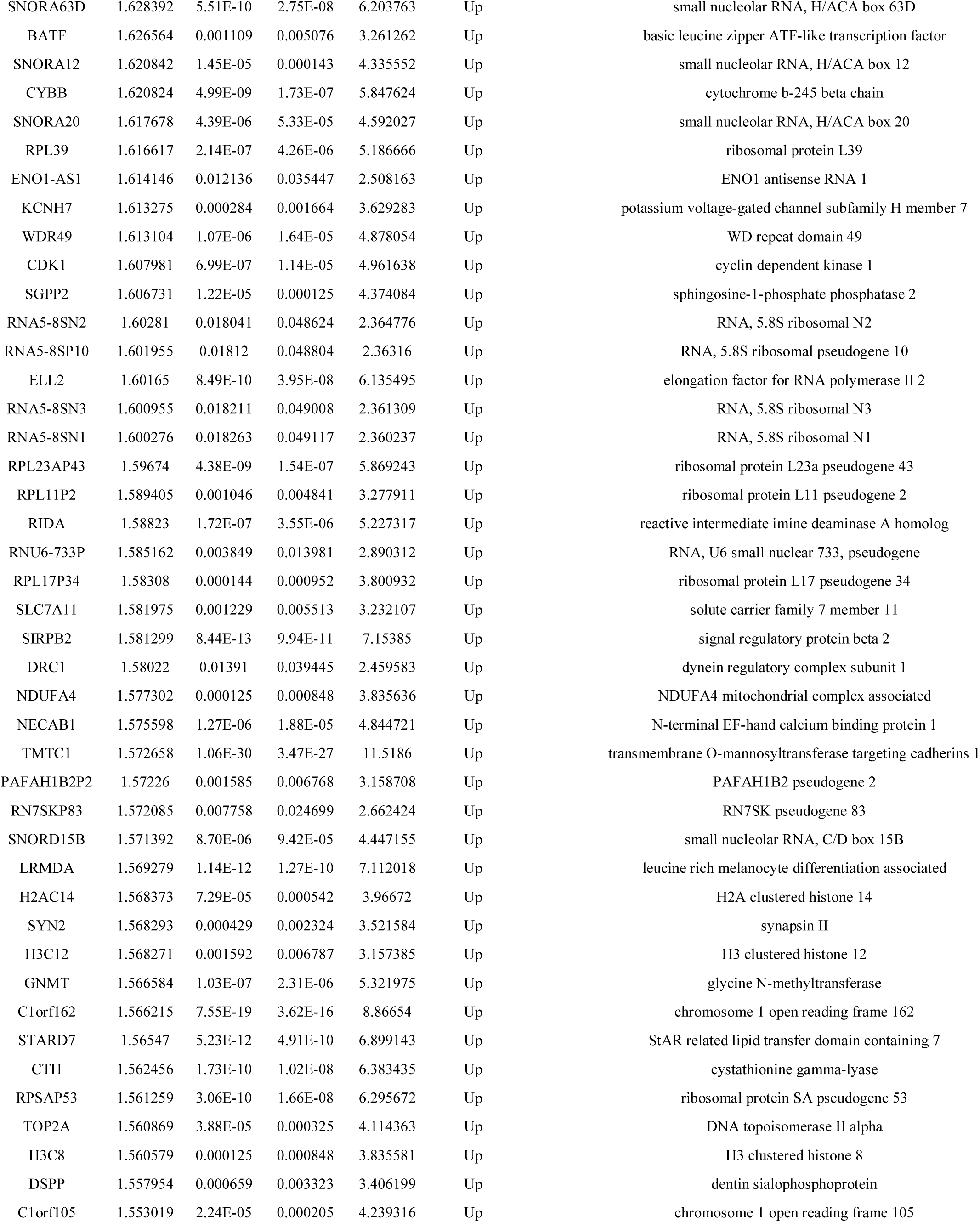
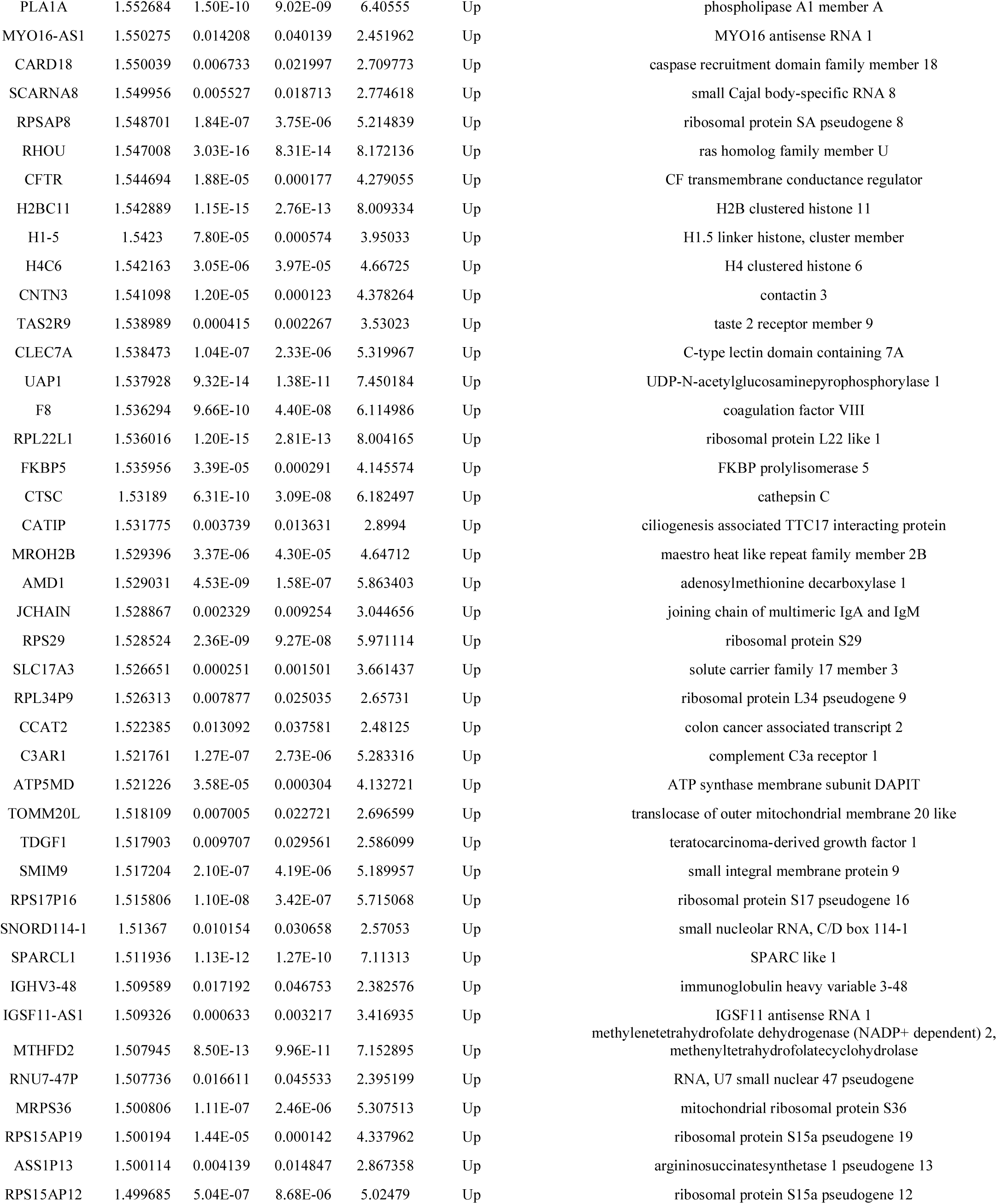
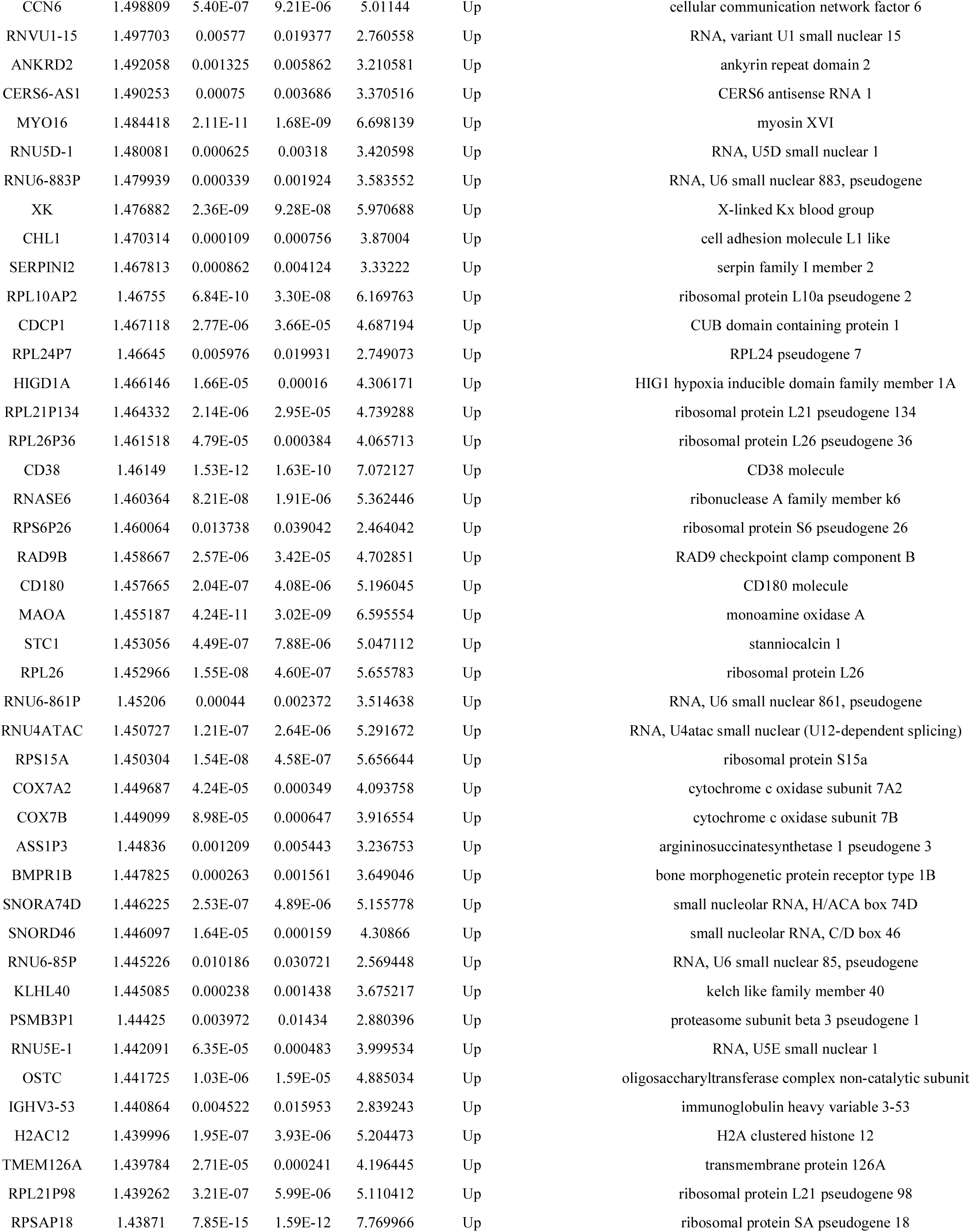
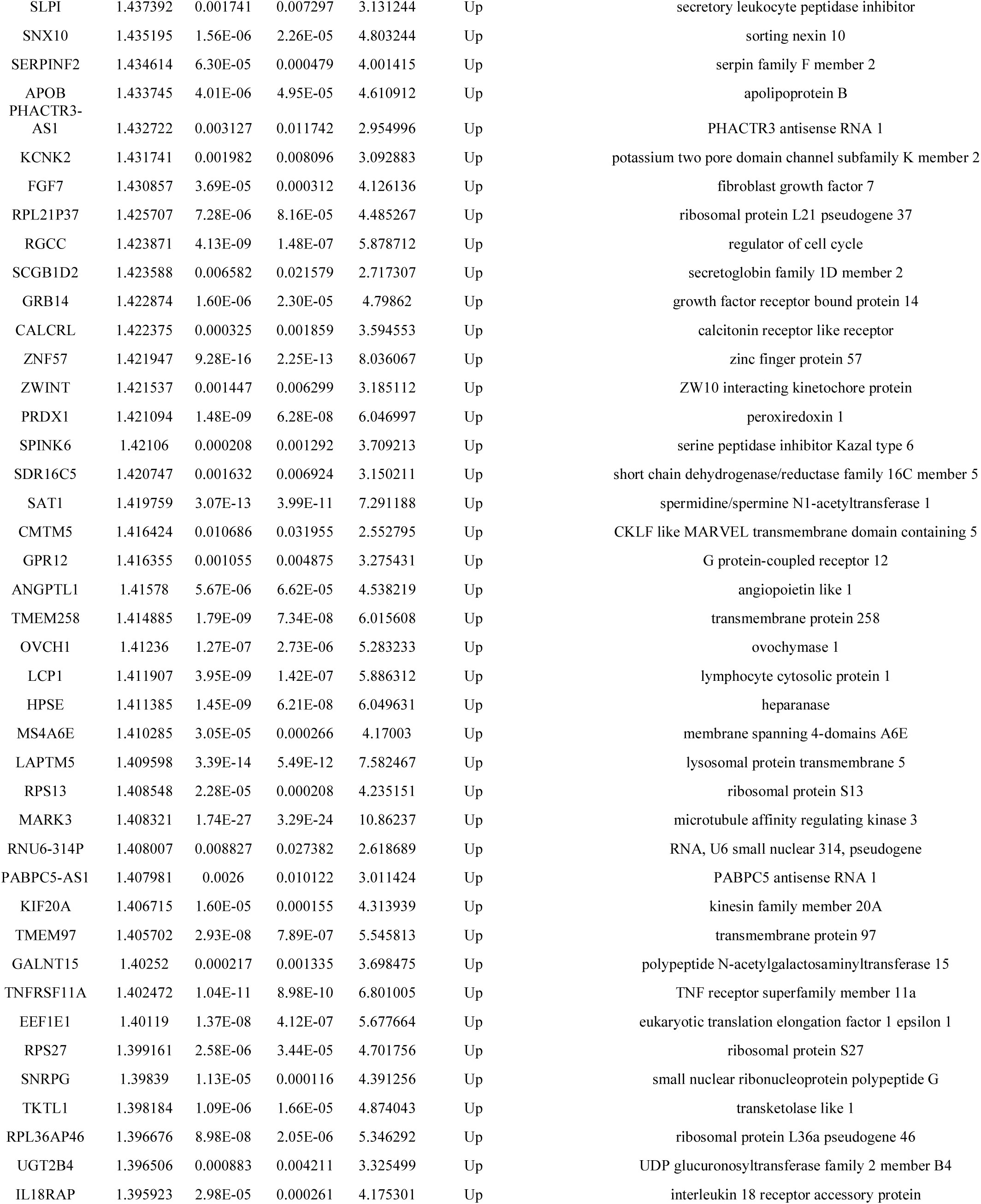
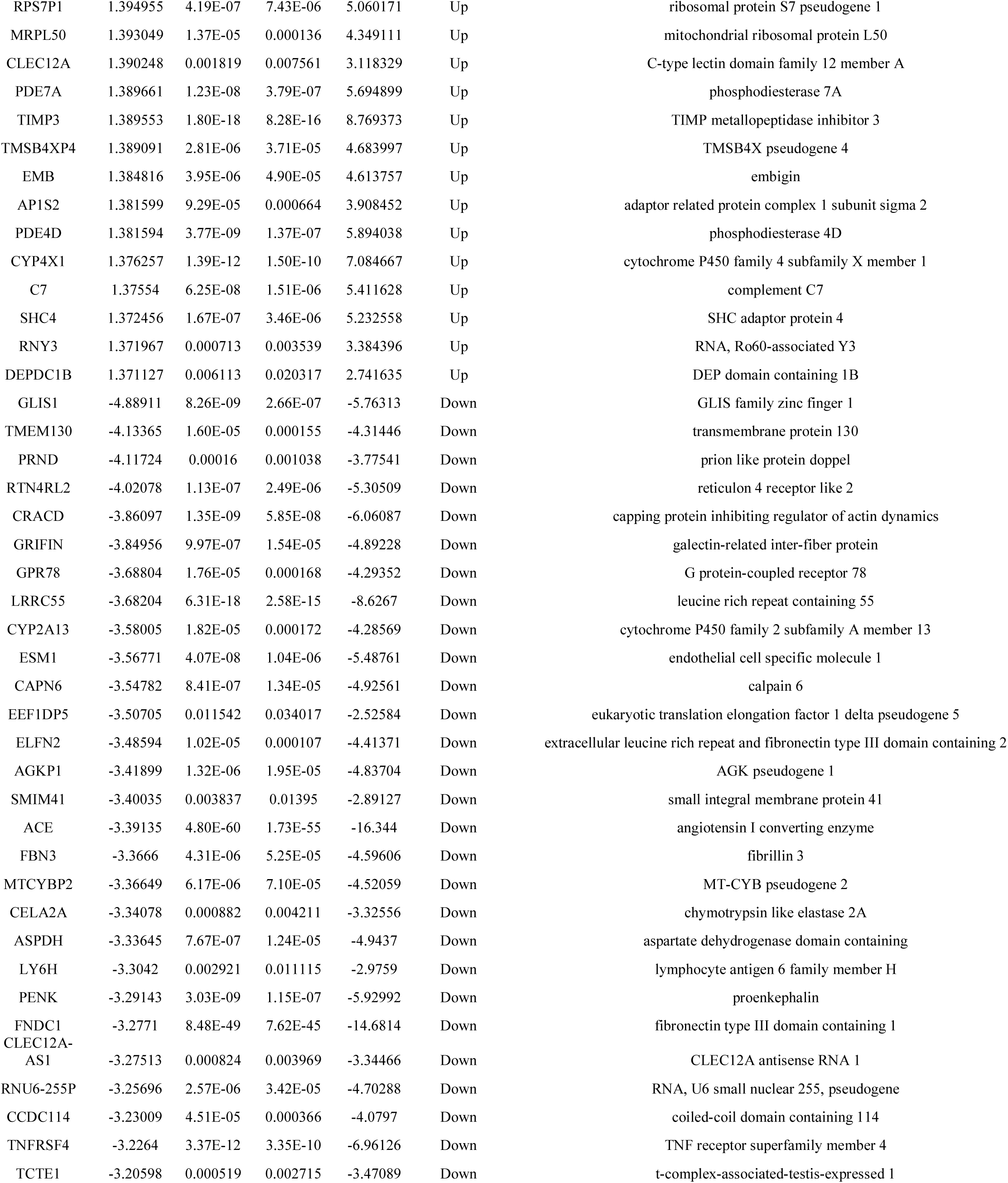
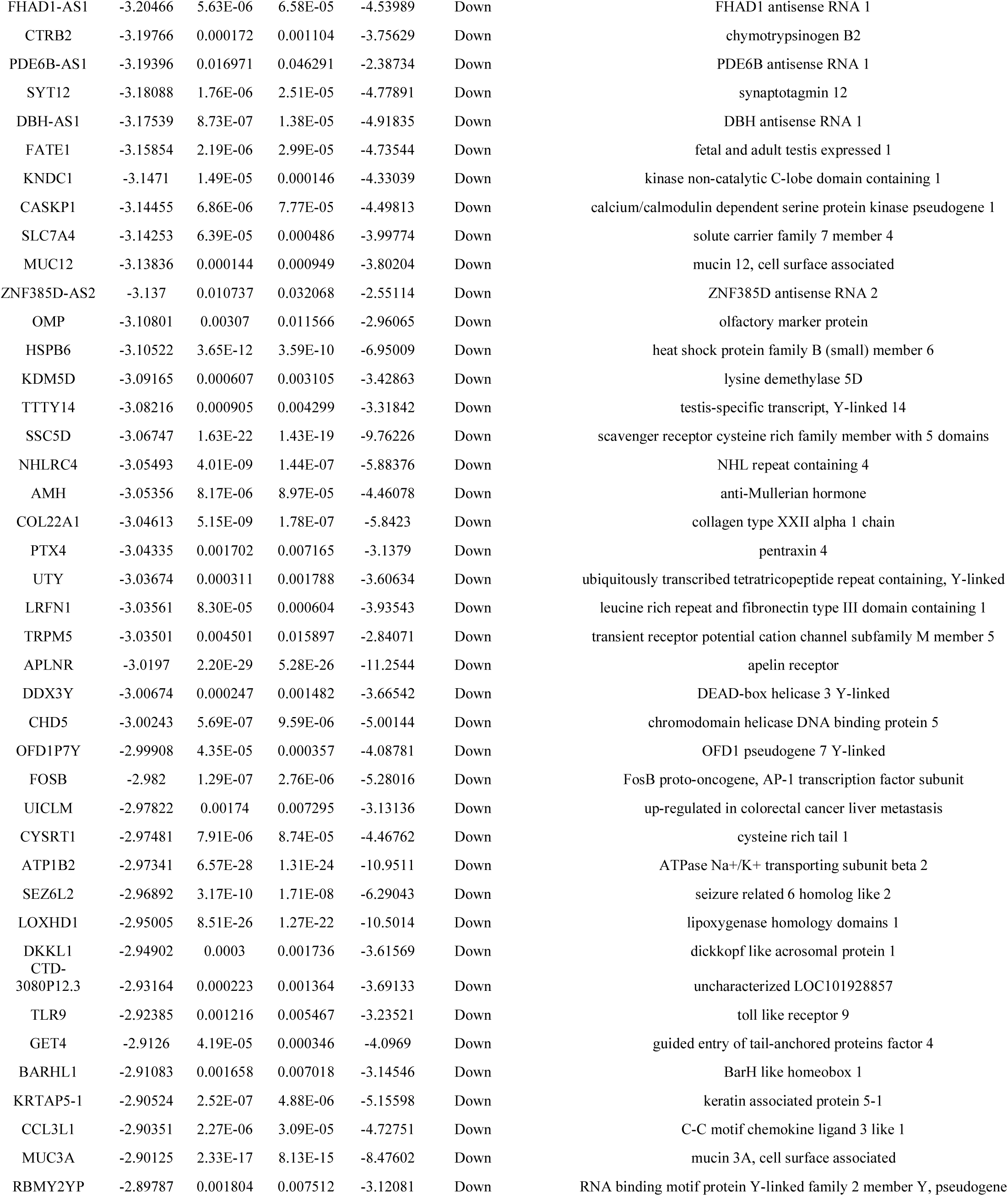
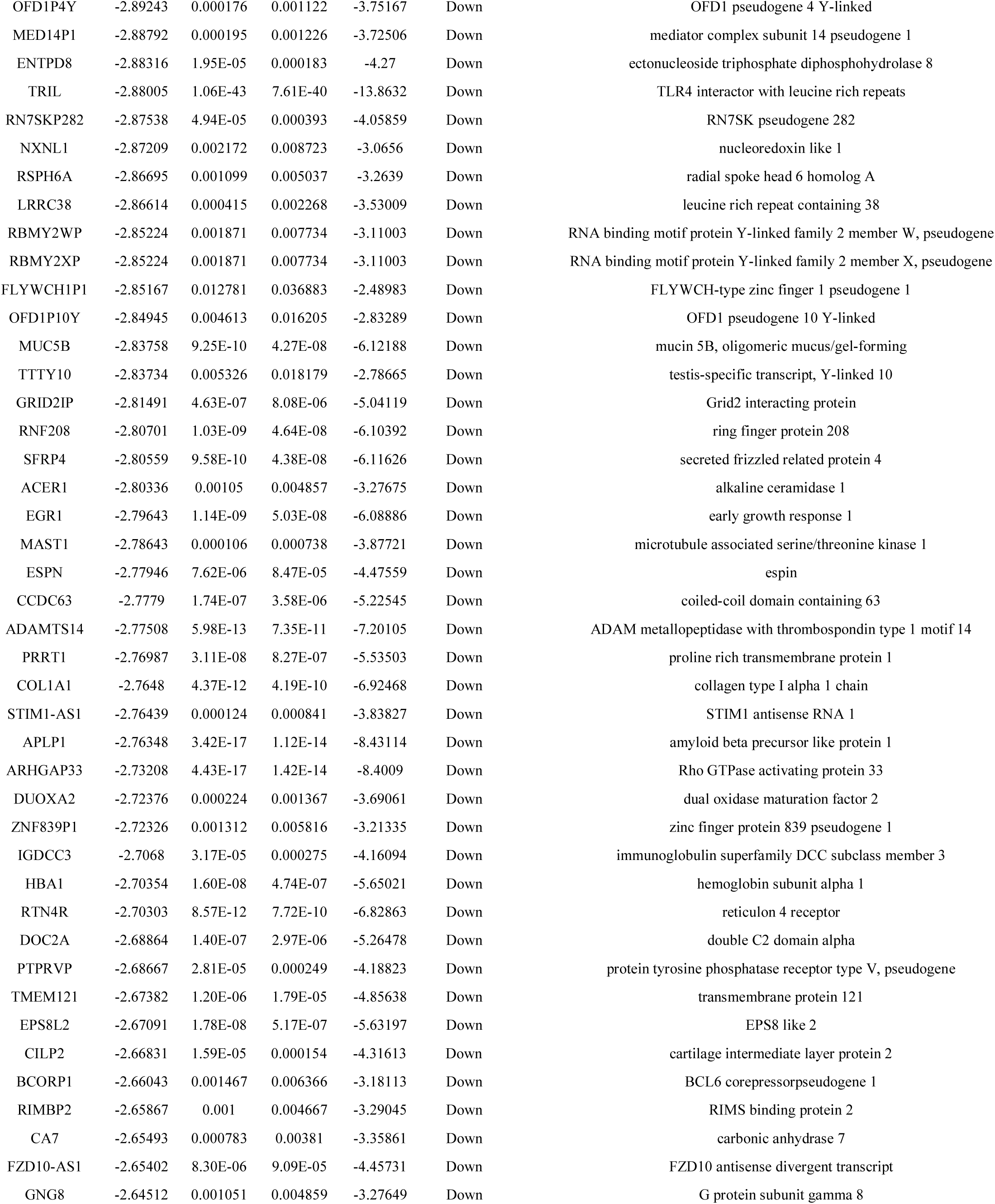
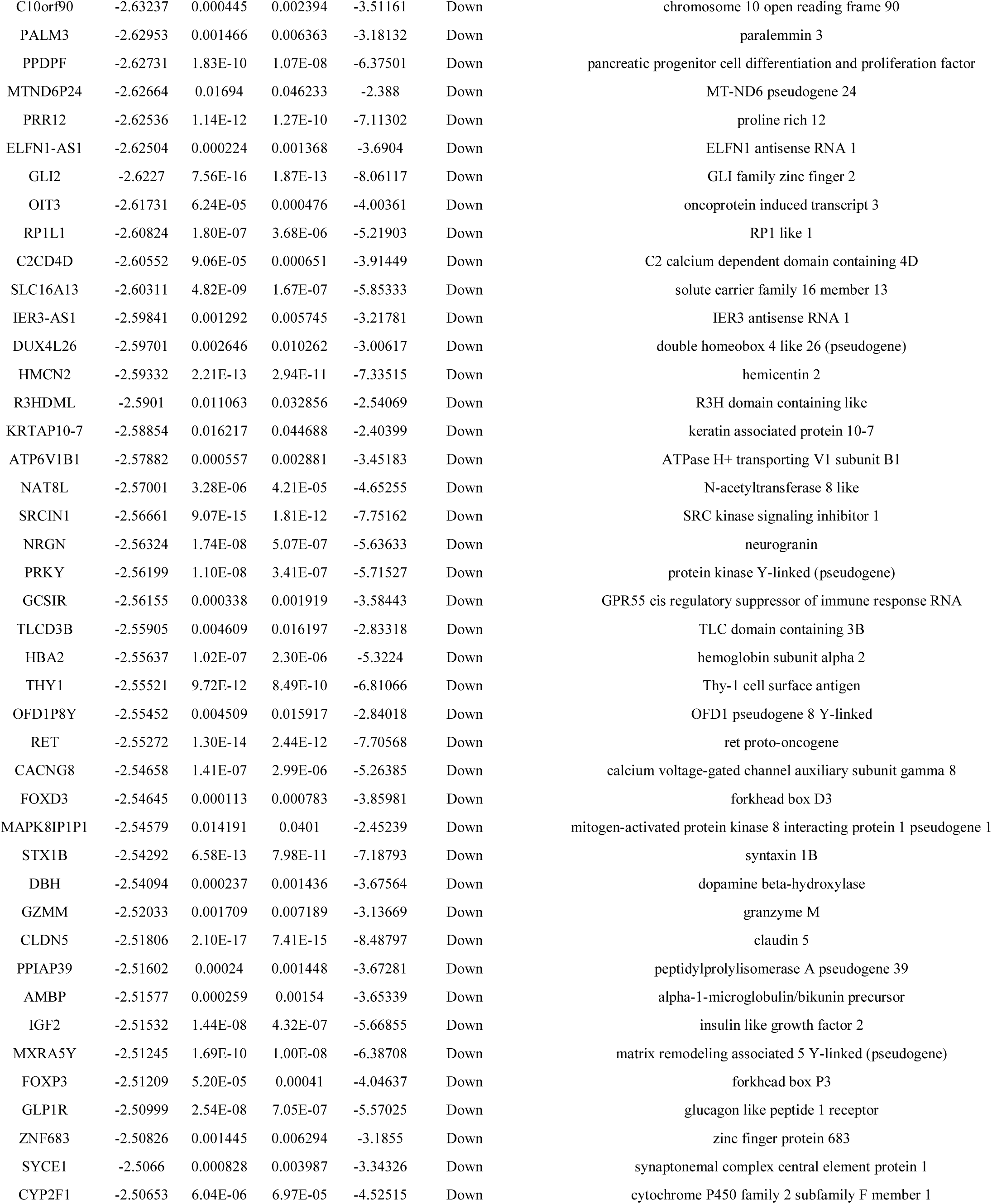
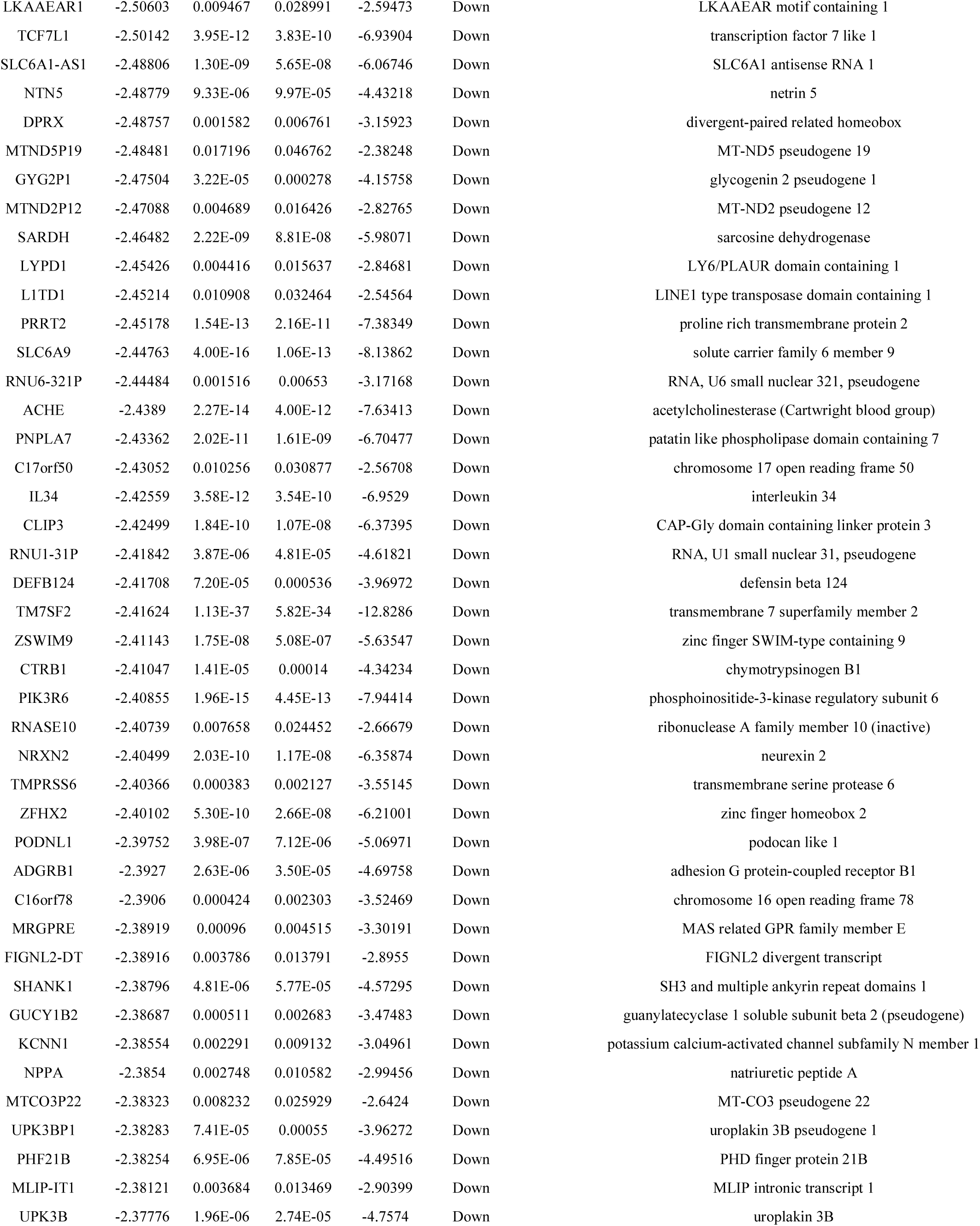
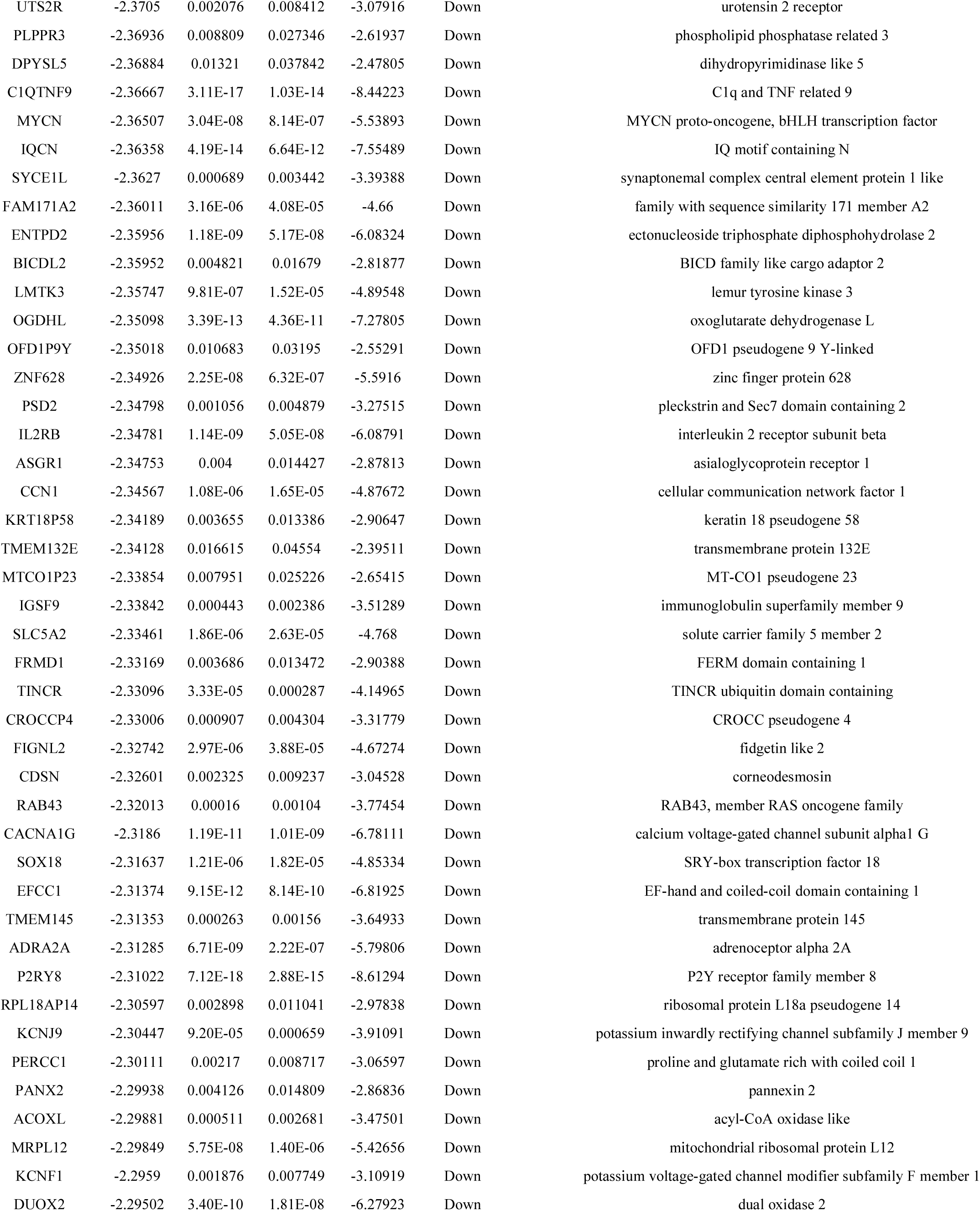
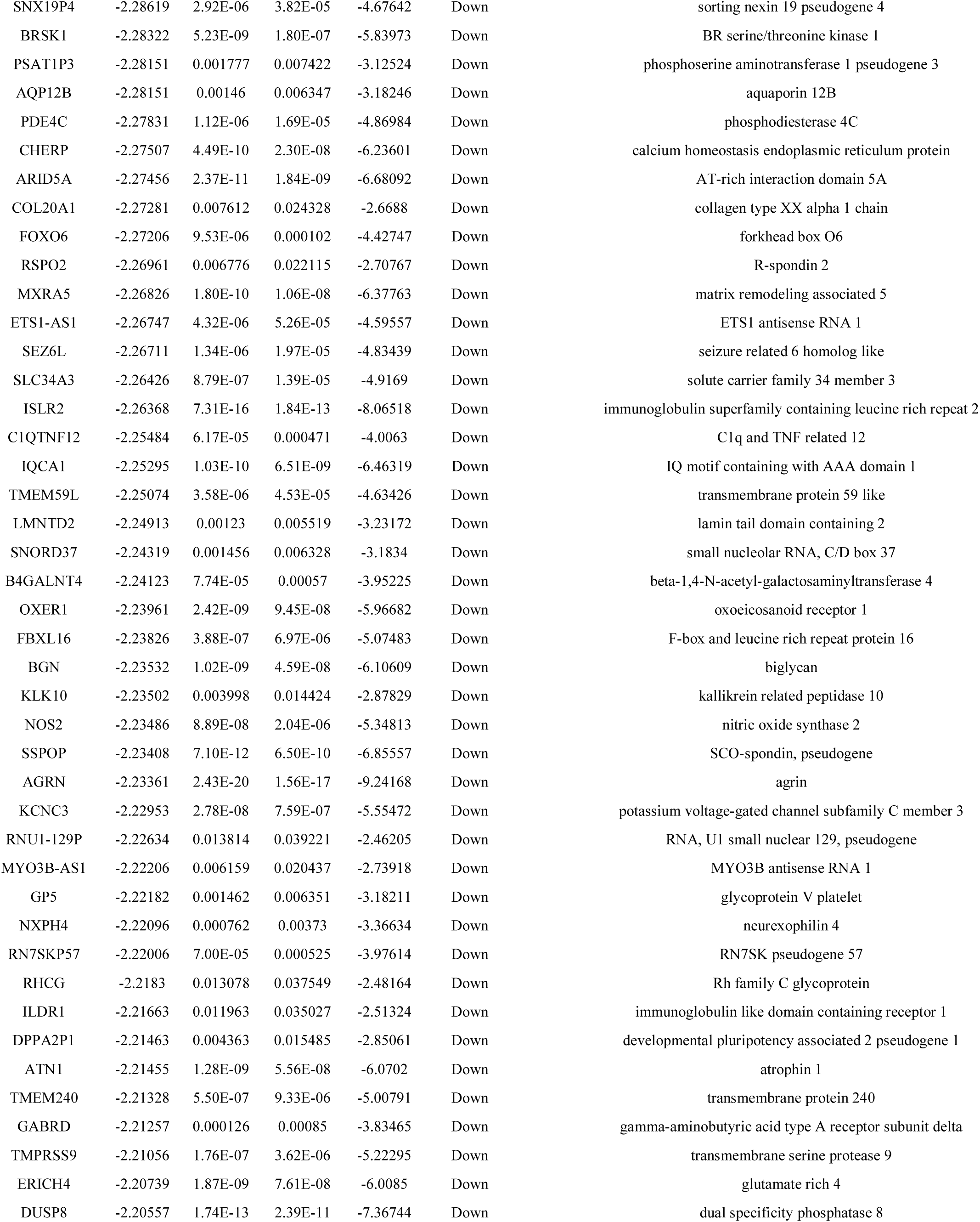
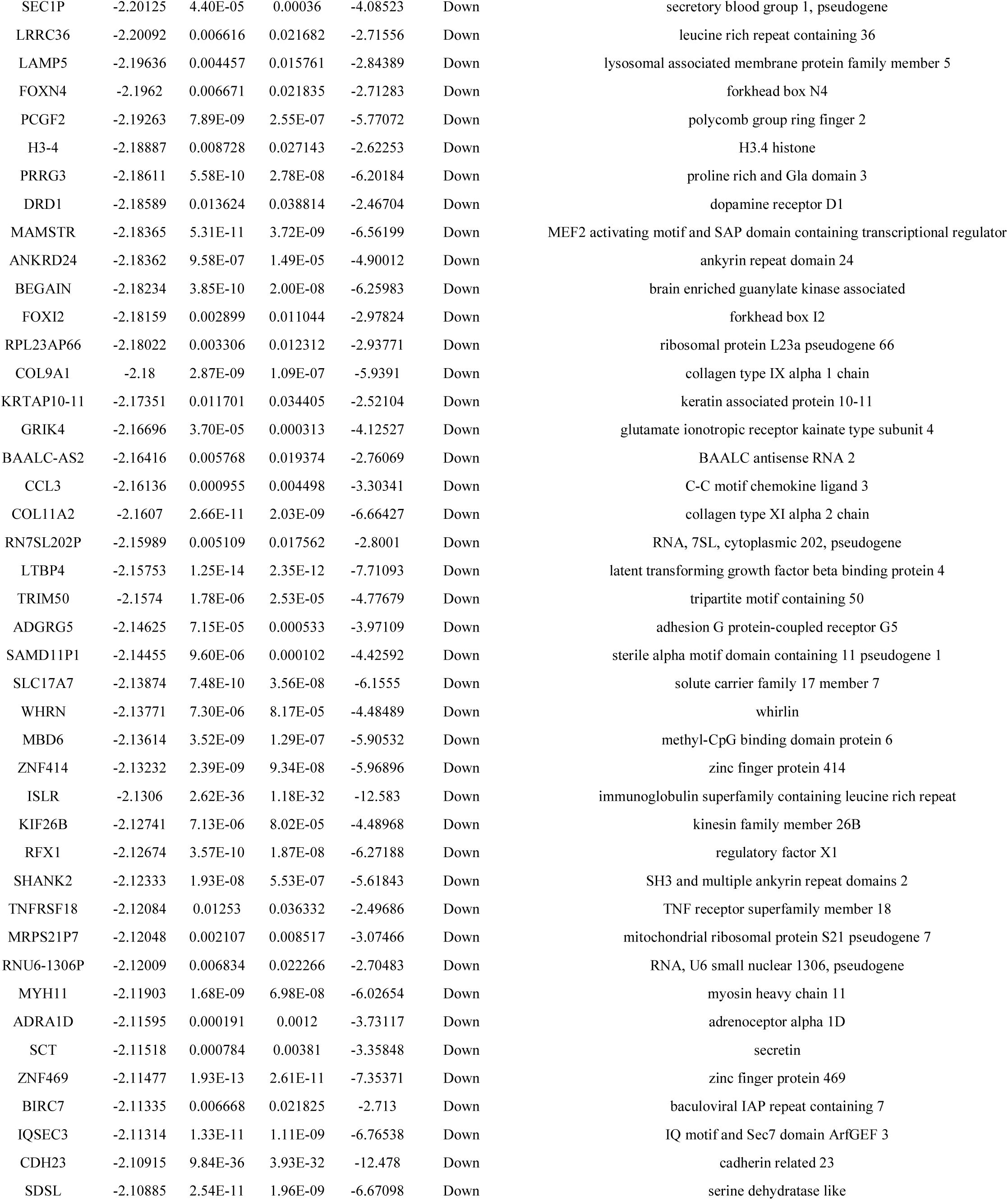
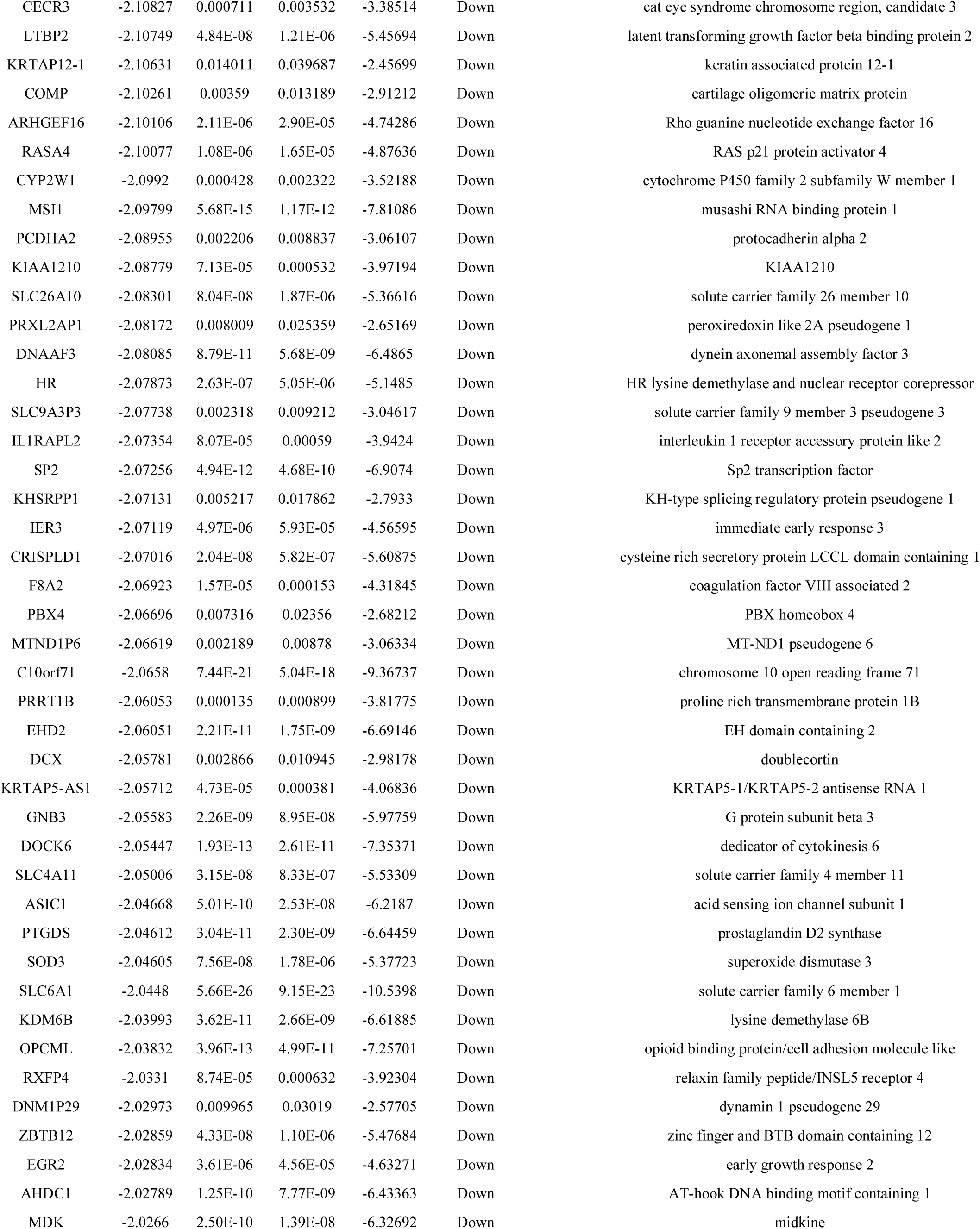
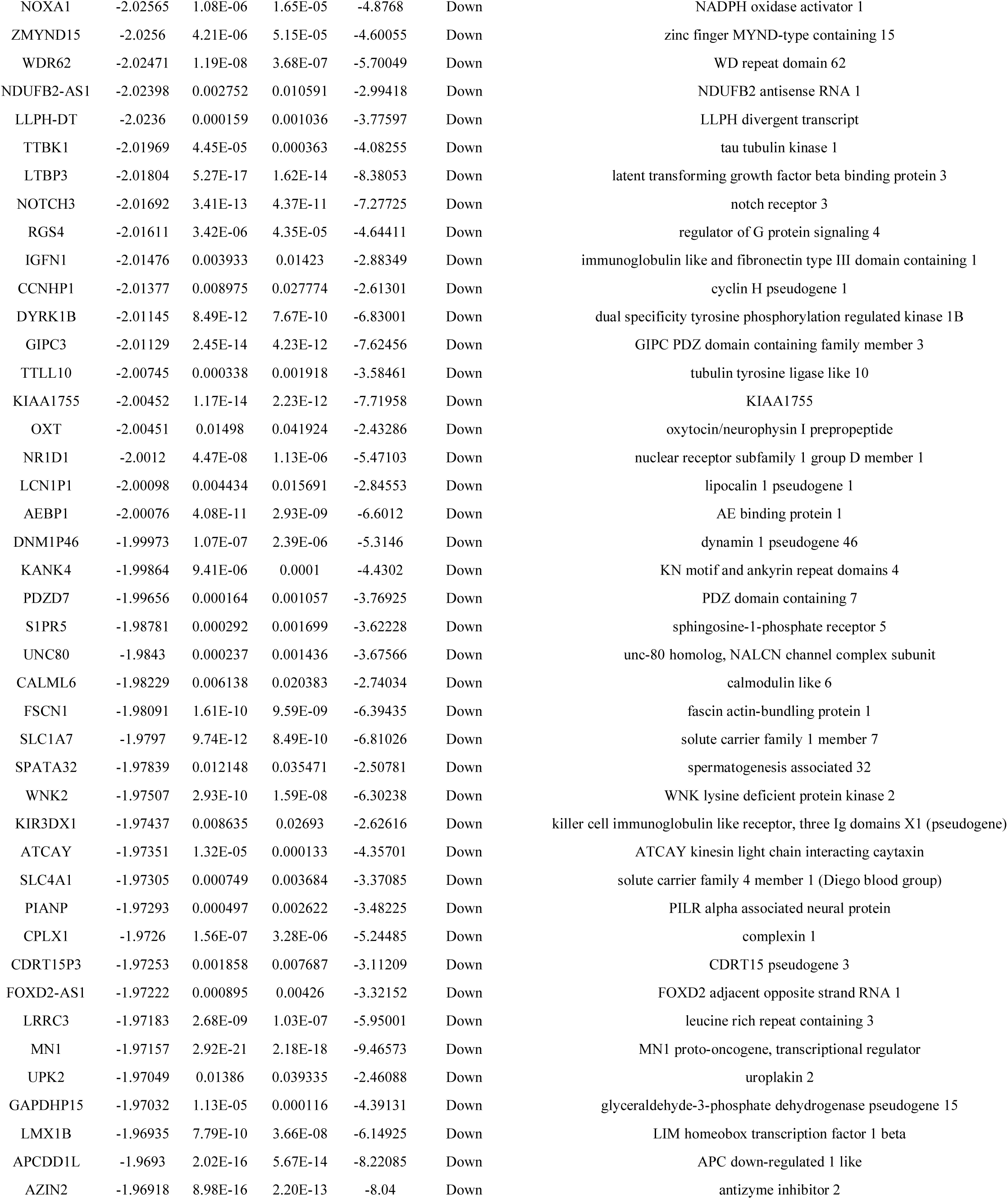
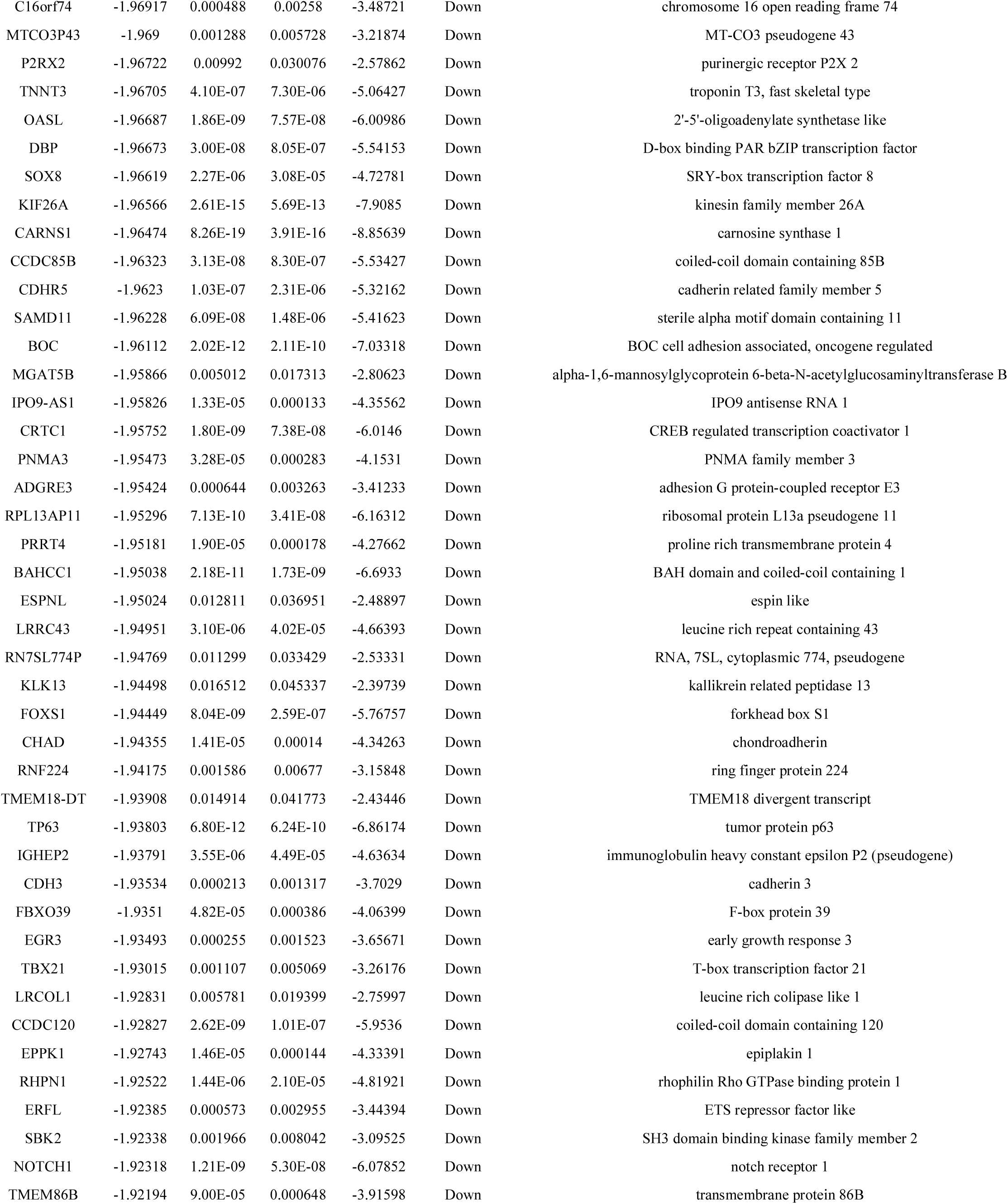
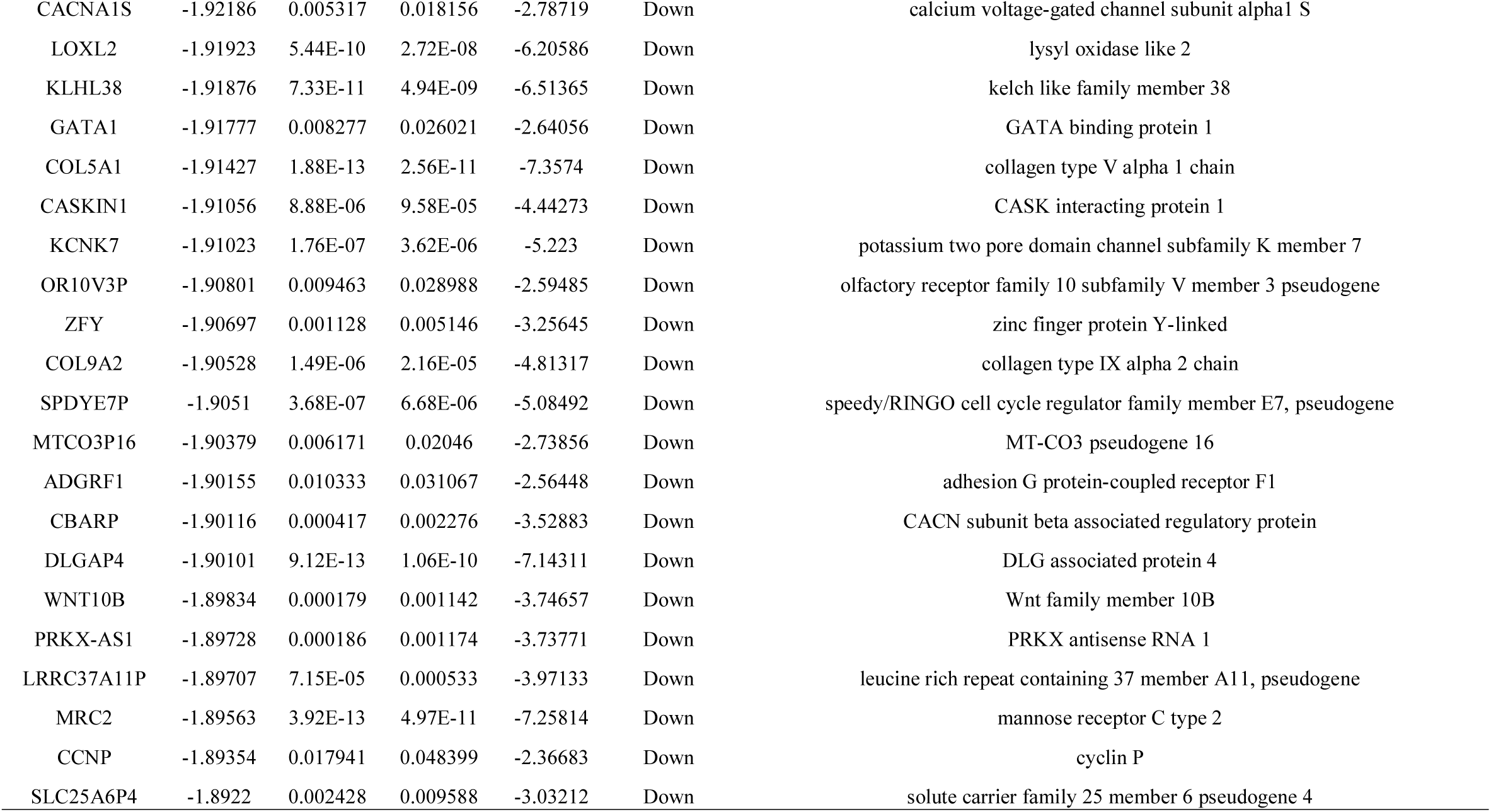
The statistical metrics for key differentially expressed genes (DEGs)

### GO and REACTOME pathway enrichment analysis of DEGs

GO analysis of genes includes BP, CC and MF. The GO enrichment analysis results showed that the up regulated genes were mainly involved in BP such as the response to stimulus and cell communication (Table 2). The down regulated genes were enriched in multicellular organismal process and anatomical structure development (Table 2). GO CC analysis indicated that the up regulated genes were mainly involved in cell periphery and membrane (Table 2). The down regulated genes were enriched in cell periphery and plasma membrane (Table 2). In addition, for the MF analysis, the up regulated genes were significantly enriched in the molecular transducer activity and transmembranesignaling receptor activity (Table 2). The down regulated genes were enriched in inorganic molecular entity transmembrane transporter activity and structural molecule activity (Table 2). REACTOME pathway enrichment analyses indicated that the up regulated genes were enriched in immune system and innate immune system (Table 3). The down regulated genes were enriched in neuronal system and extracellular matrix organization (Table 3).

**Table 2.**
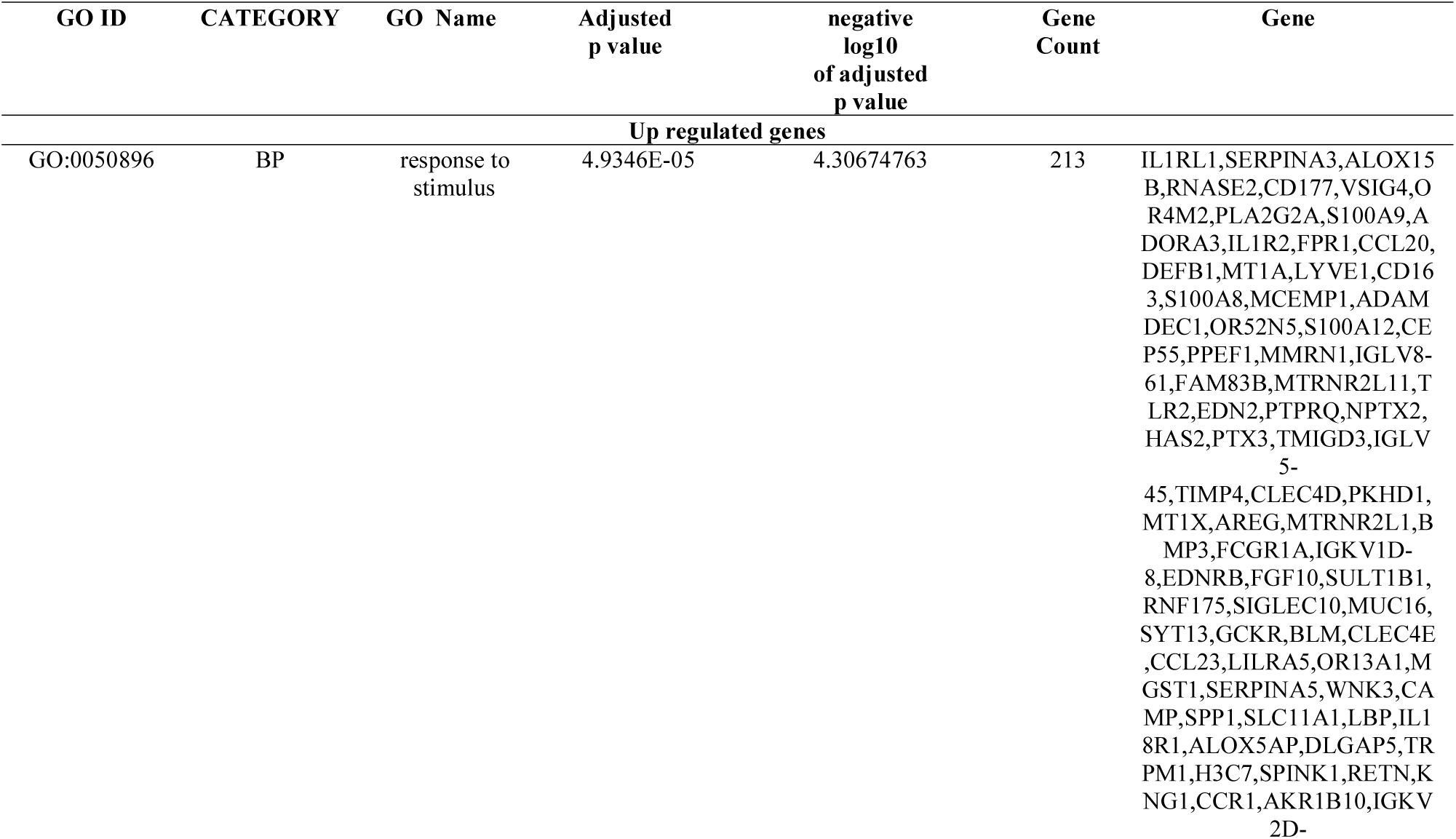
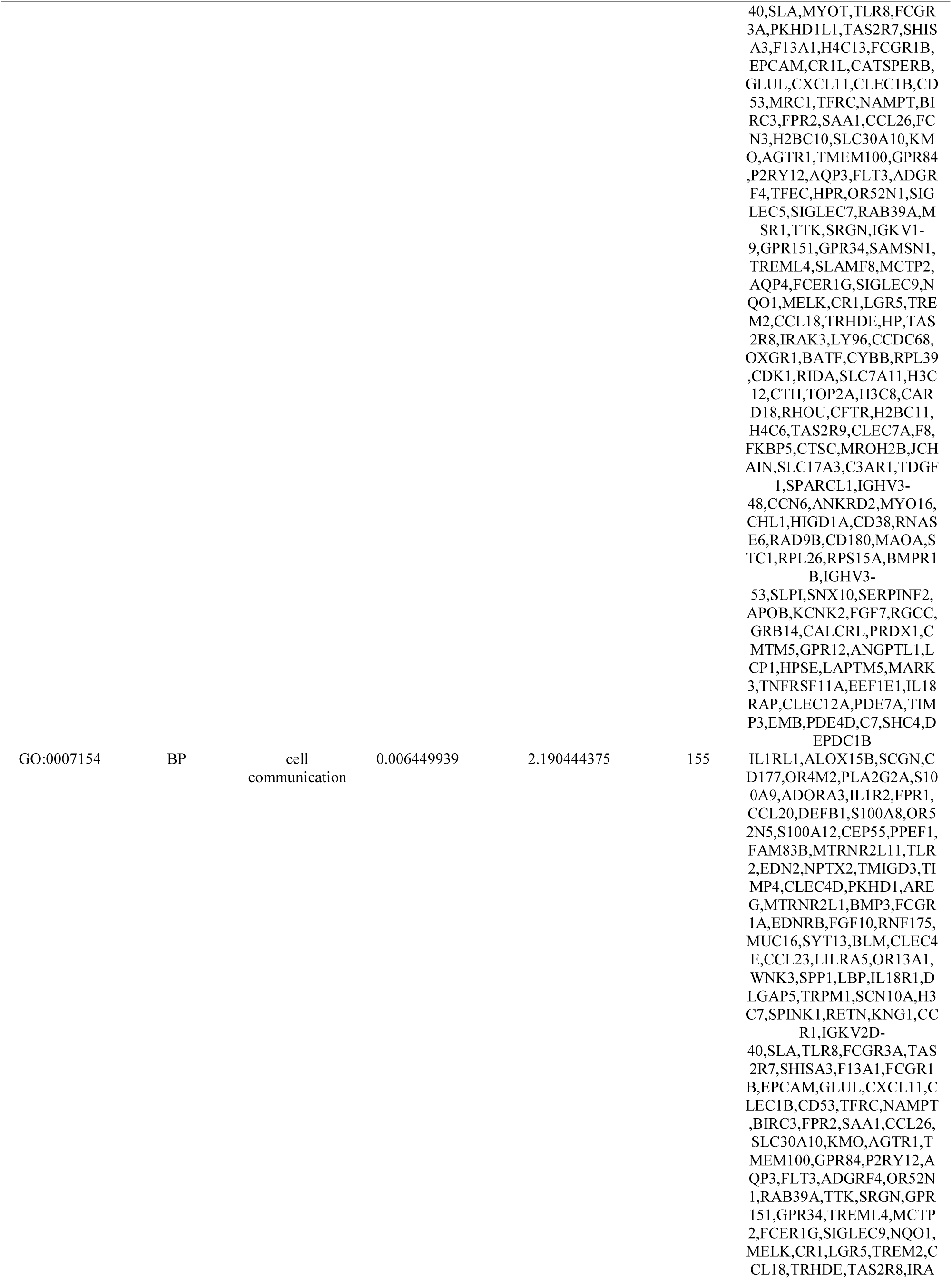
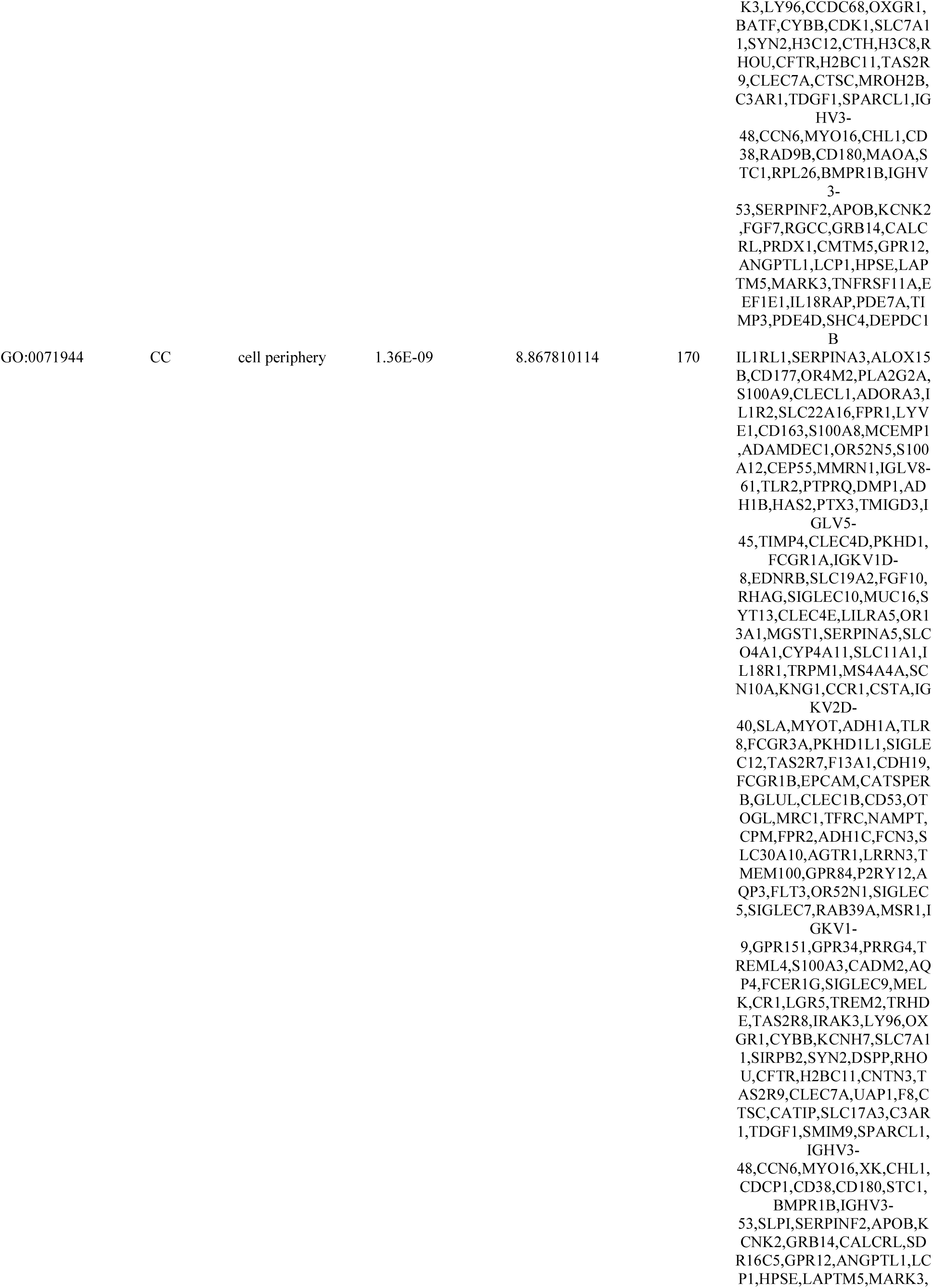
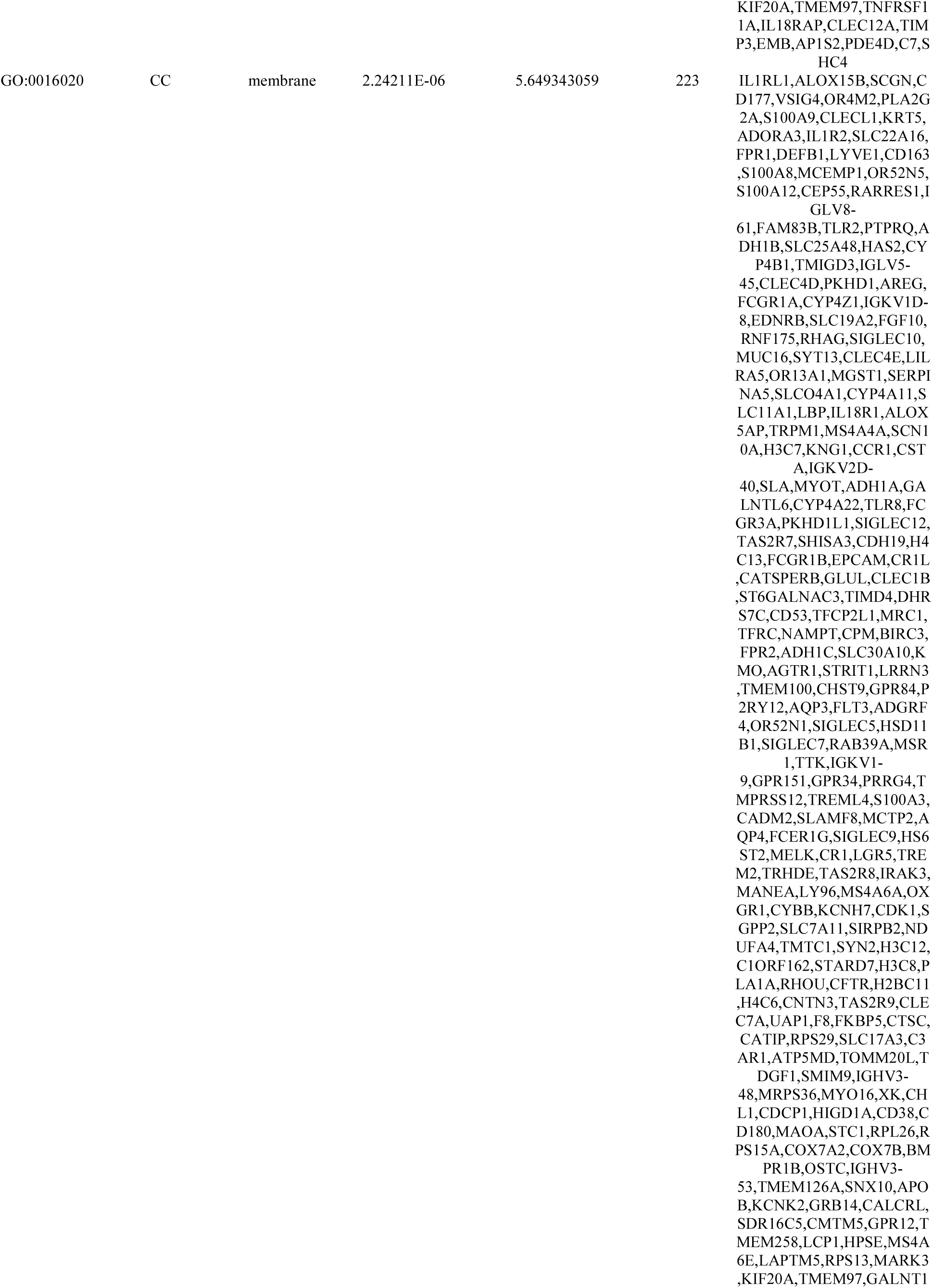
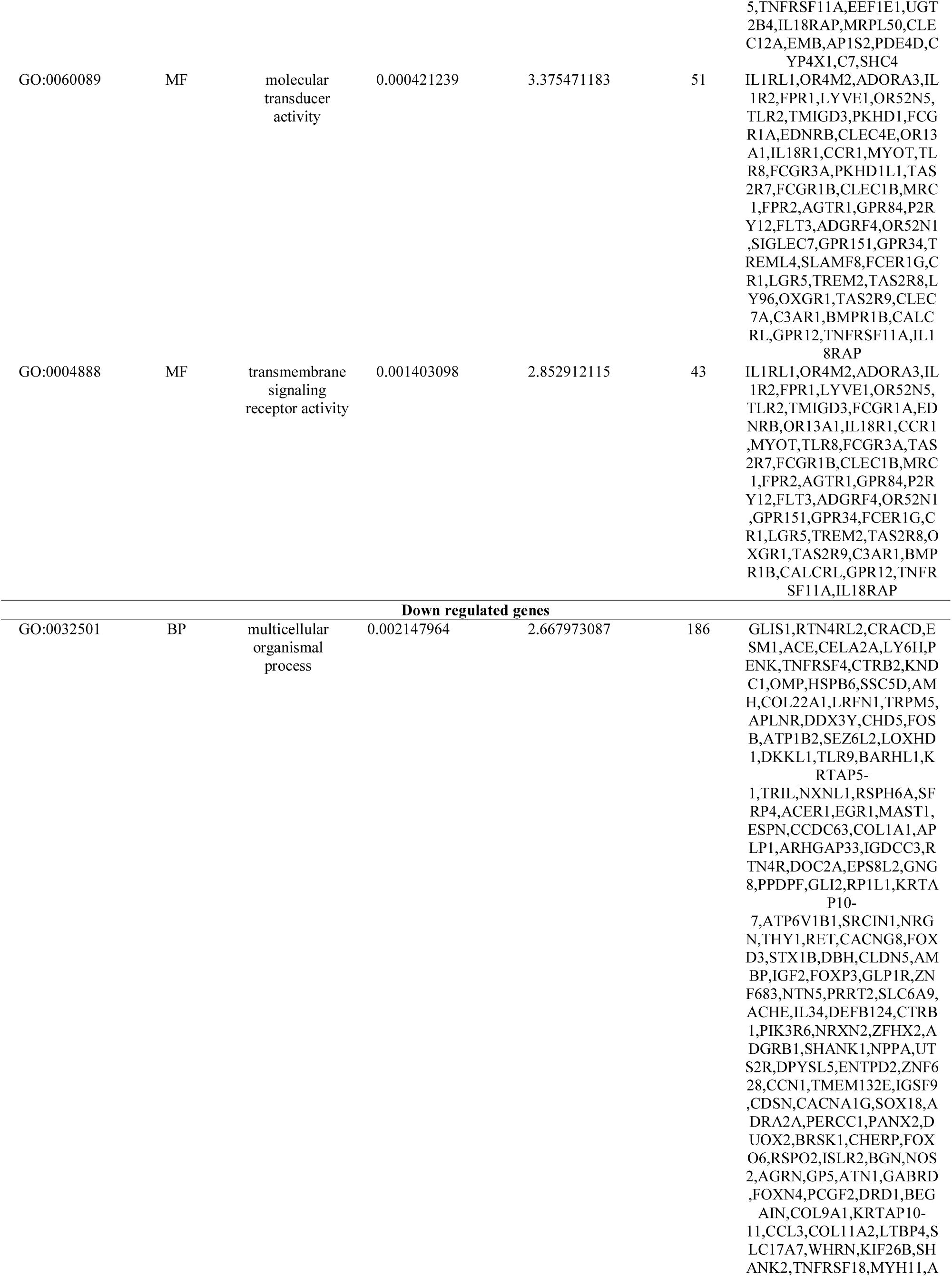
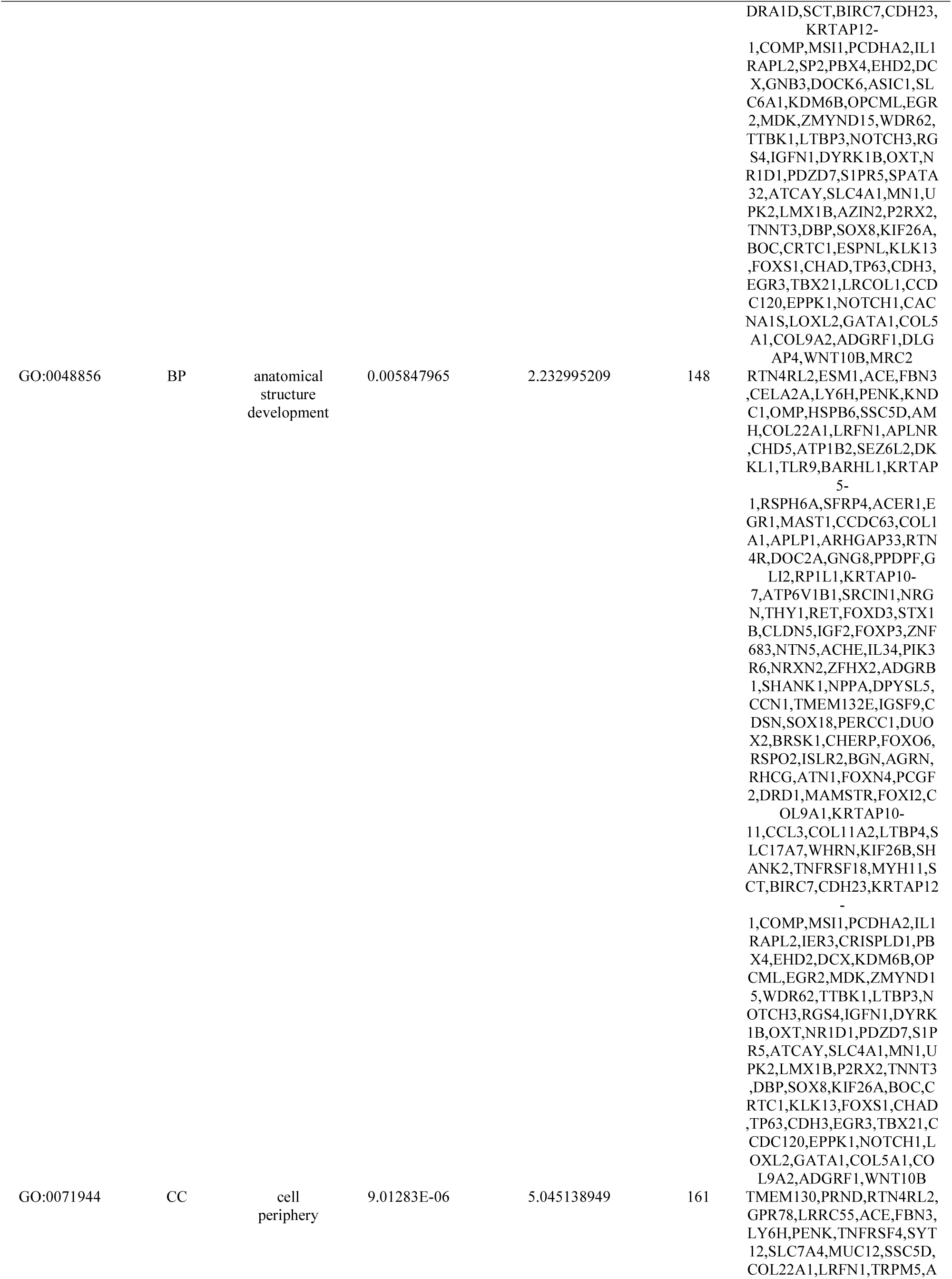
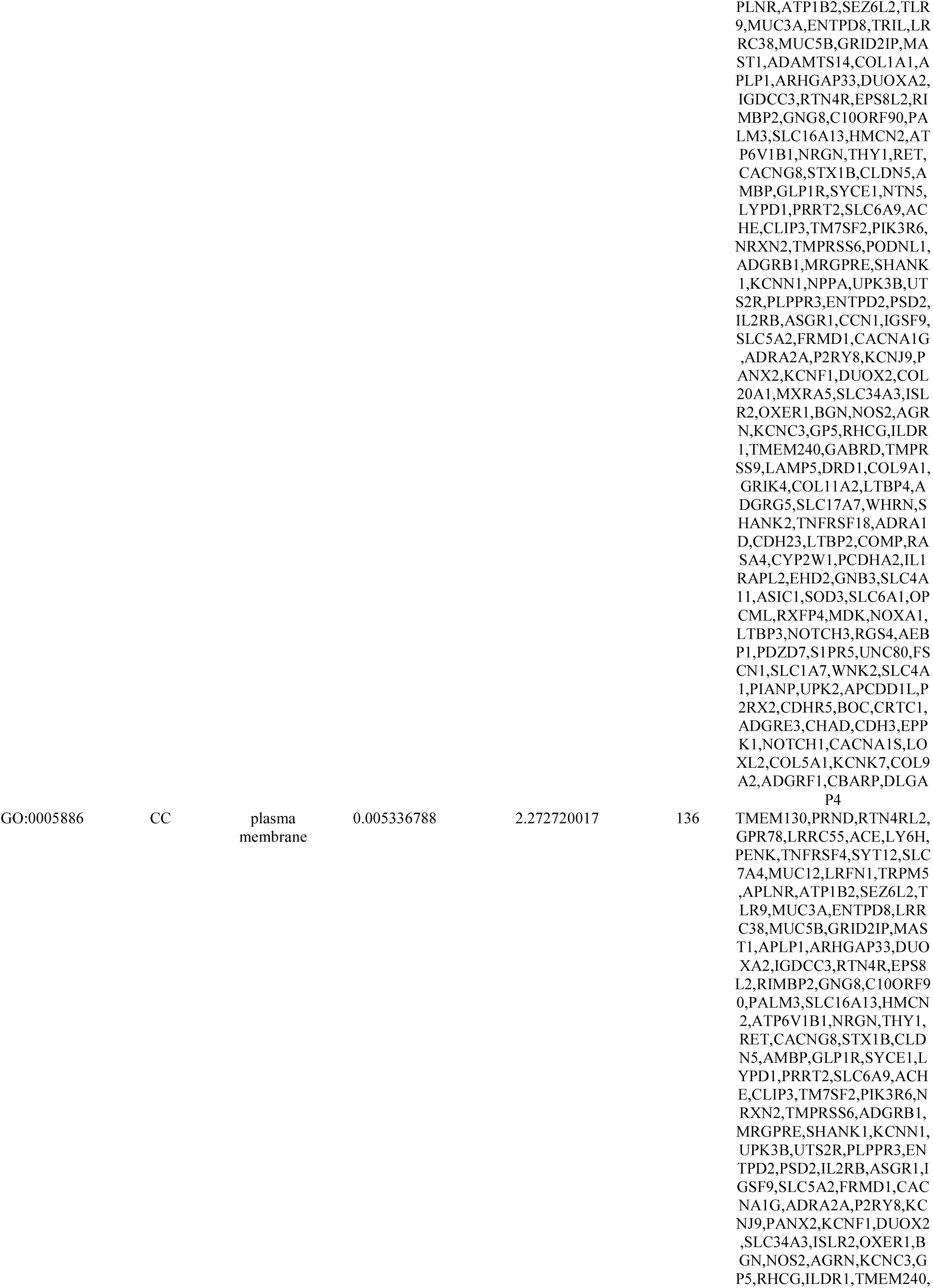
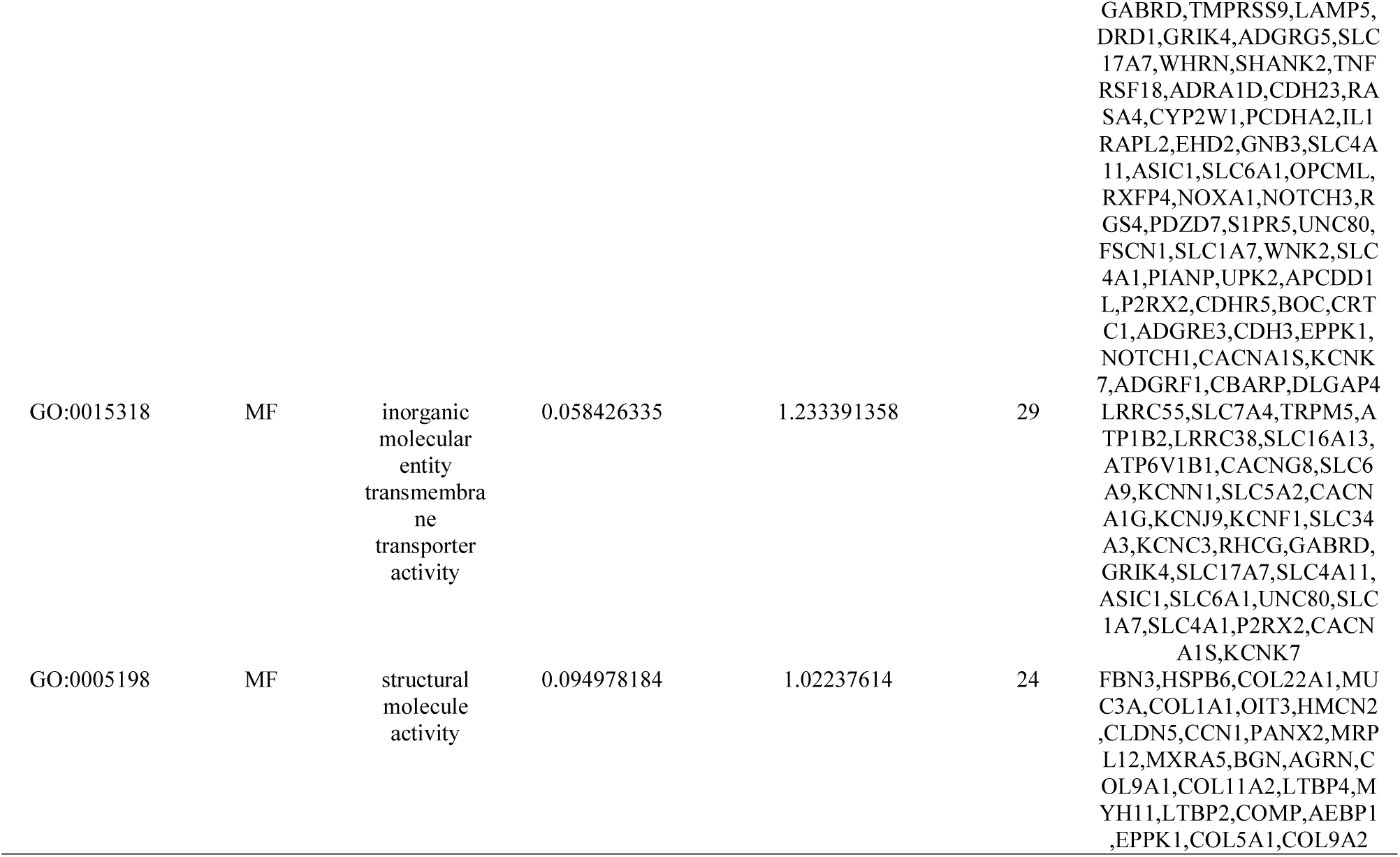
The enriched GO terms of the up and down regulated differentially expressed genes

**Table 3.**
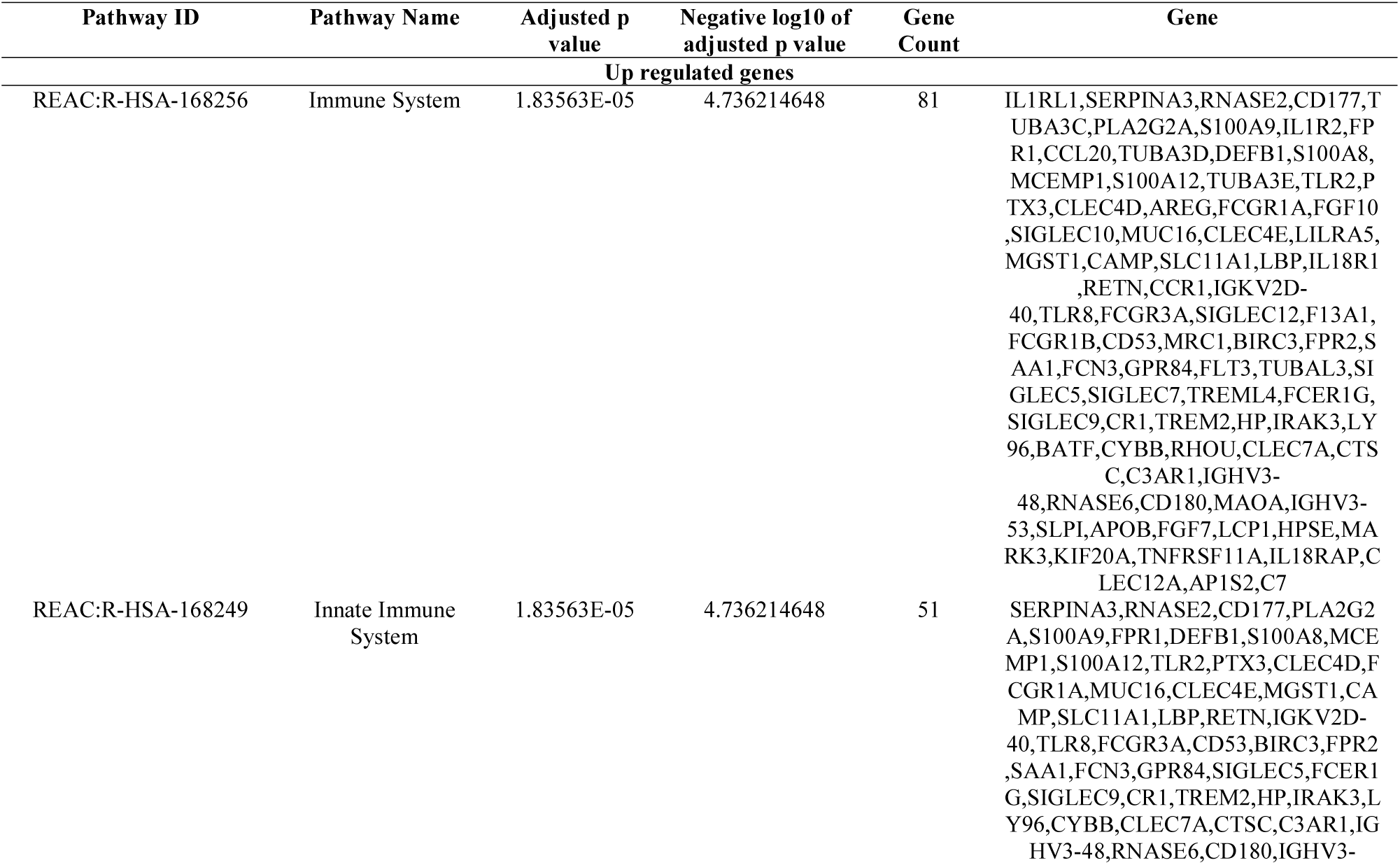
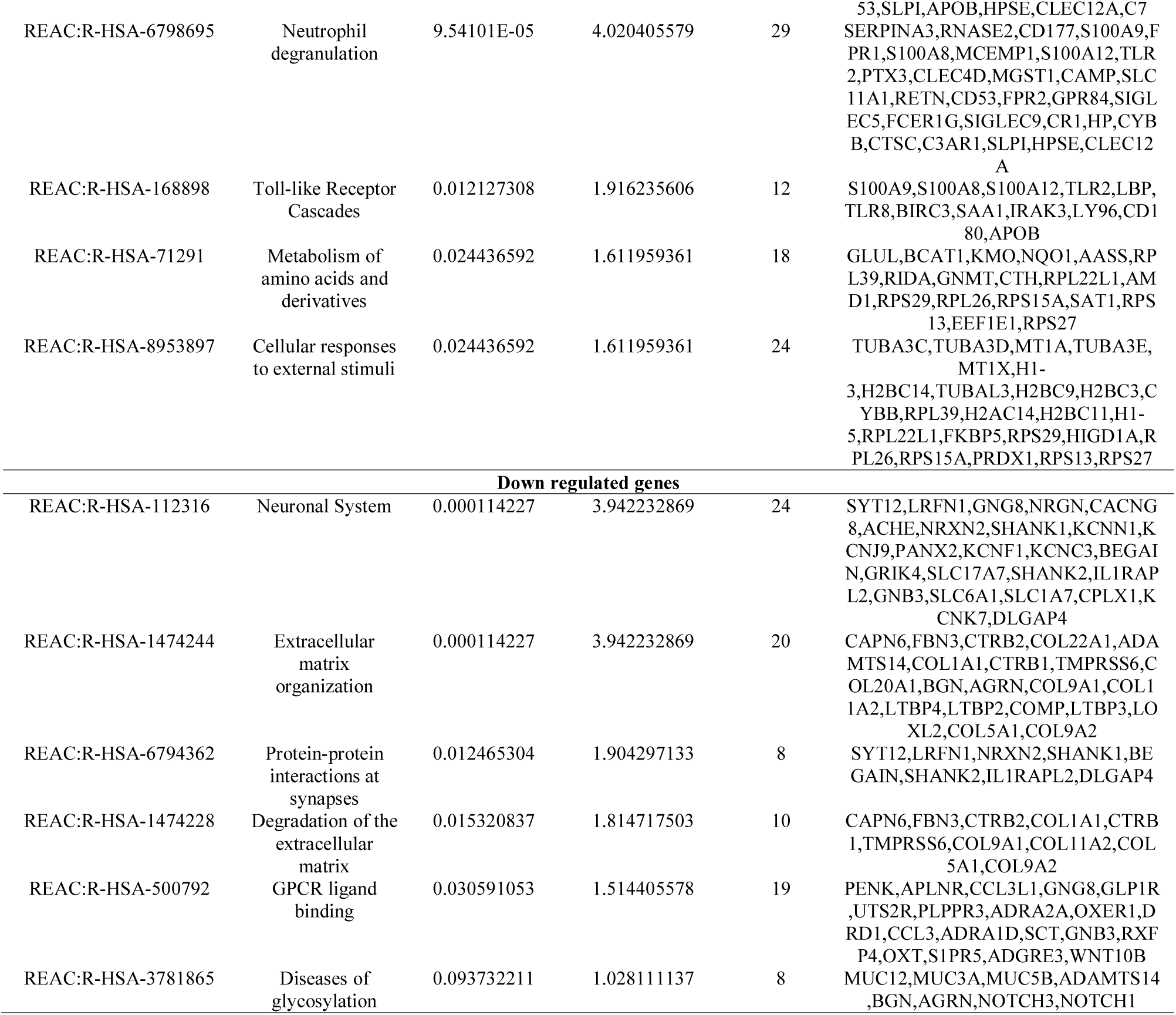
The enriched pathway terms of the up and down regulated differentially expressed genes

### Construction of the PPI network and module analysis

The PPI network of DEGs consisting of 6292 nodes and 12582 edges was constructed in the HiPPIE database. Then it was visualized through Cytoscape (Fig. 3). Based on the HiPPIE database, the DEGs with the highest PPI scores identified by the 4 centrality methods are shown in Table 4. After repeated genes removing, the hub genes were obtained using the 4 centrality methods, including CFTR, CDK1, RPS13, RPS15A, RPS27, NOTCH1, MRPL12, NOS2, CCDC85B and ATN1. A significant modules were constructed from the PPI network of the DEGs using PEWCC1, including module 1 had 59 nodes and 203 edges (Fig. 4A) and module 2 had 21 nodes and 41 edges (Fig. 4B). Enrichment analysis demonstrated that modules 1 and 2 may be associated with metabolism of amino acids and derivatives, cellular responses to external stimuli, response to stimulus, membrane, diseases of glycosylation and multicellular organismal process.

**Fig. 3.**
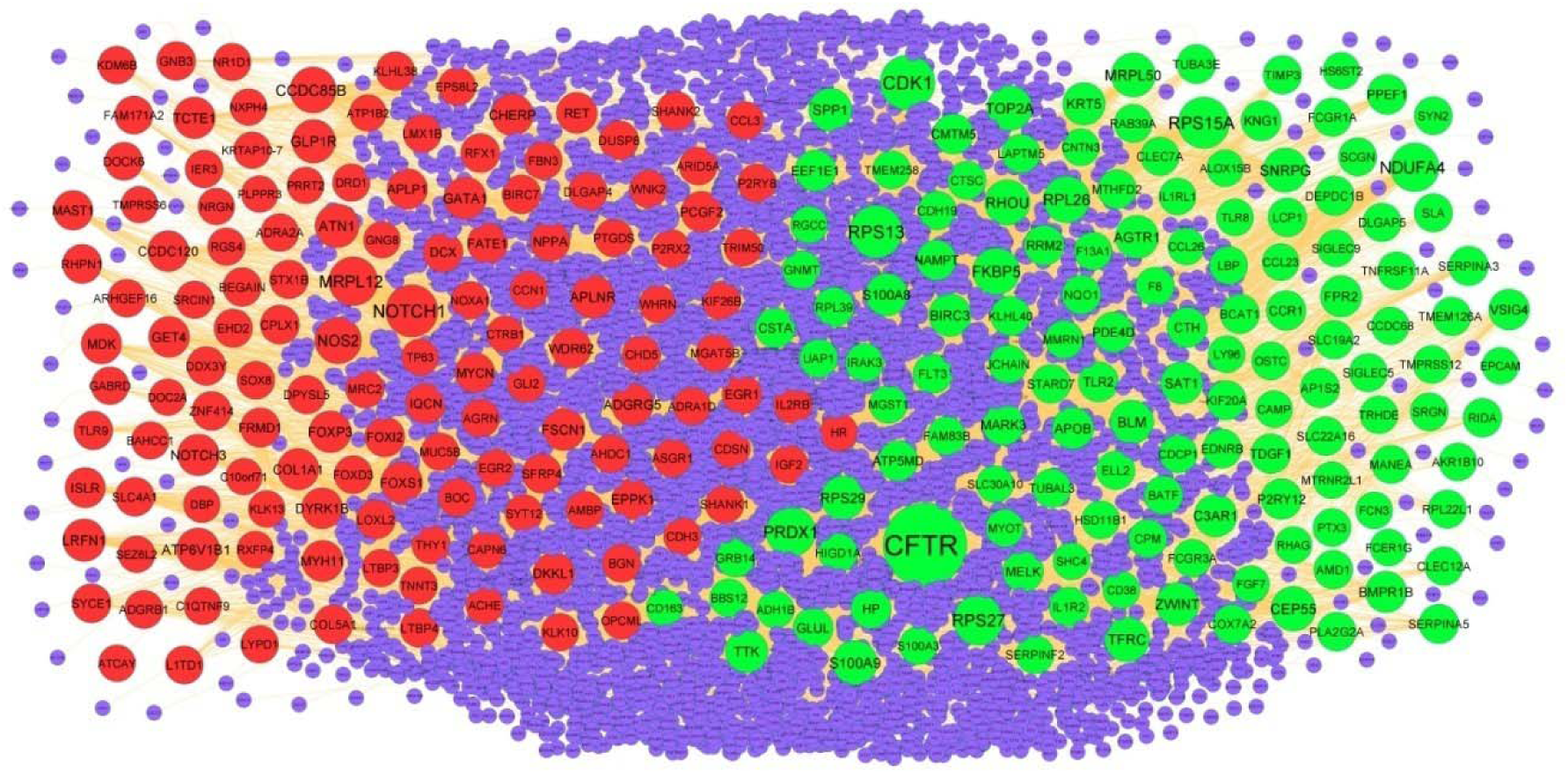
PPI network of DEGs. Up regulated genes are marked in green; down regulated genes are marked in red

**Fig. 4.**
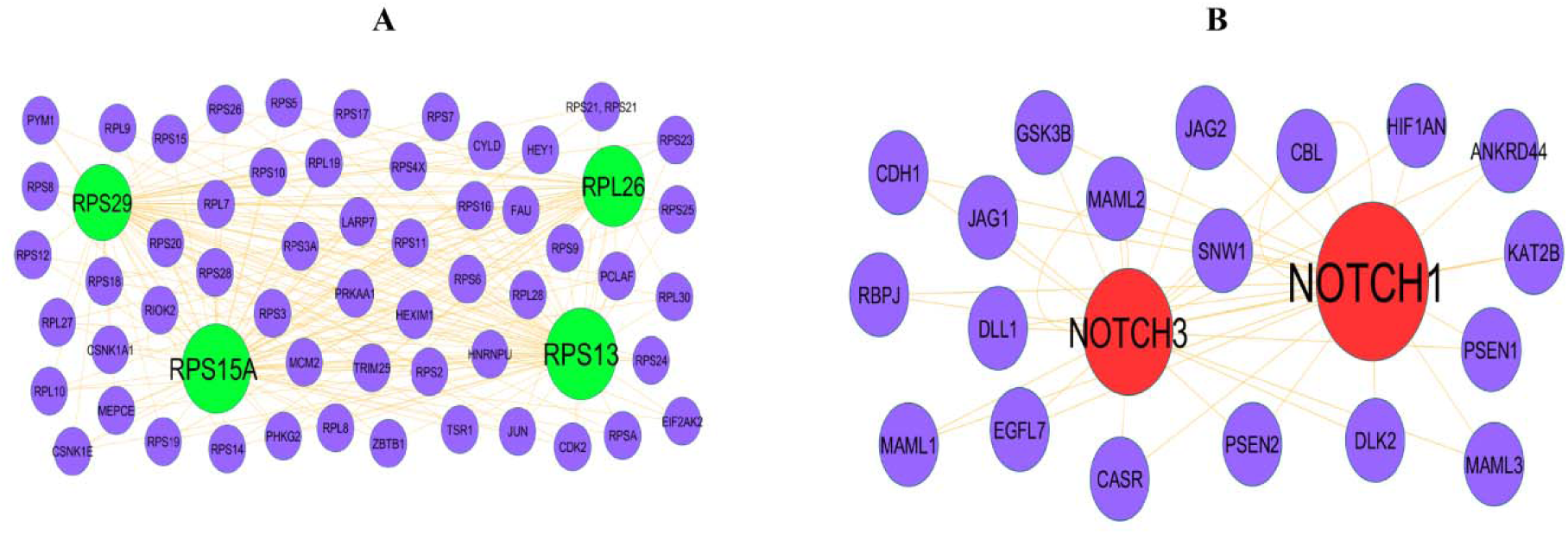
Modules of isolated form PPI of DEGs. (A) The most significant module was obtained from PPI network with 59 nodes and 203 edges for up regulated genes (B) The most significant module was obtained from PPI network with 21 nodes and 41 edges for down regulated genes. Up regulated genes are marked in green; down regulated genes are marked in red

**Table 4.**
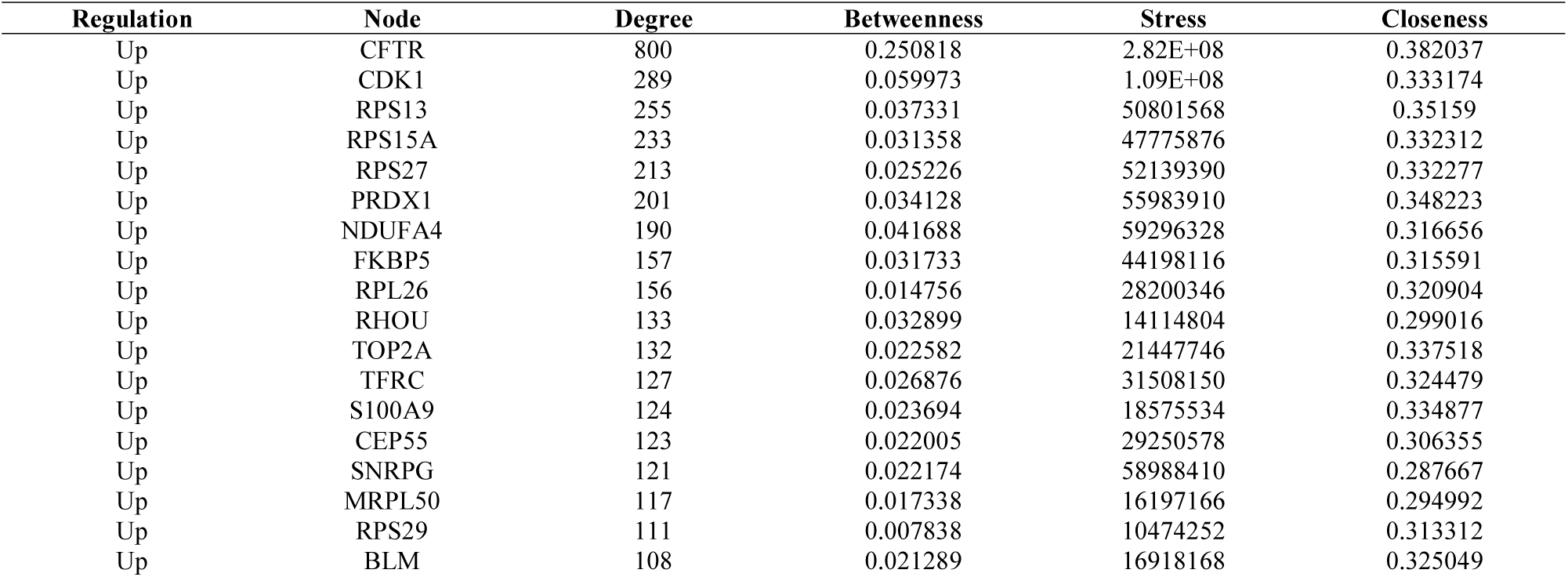
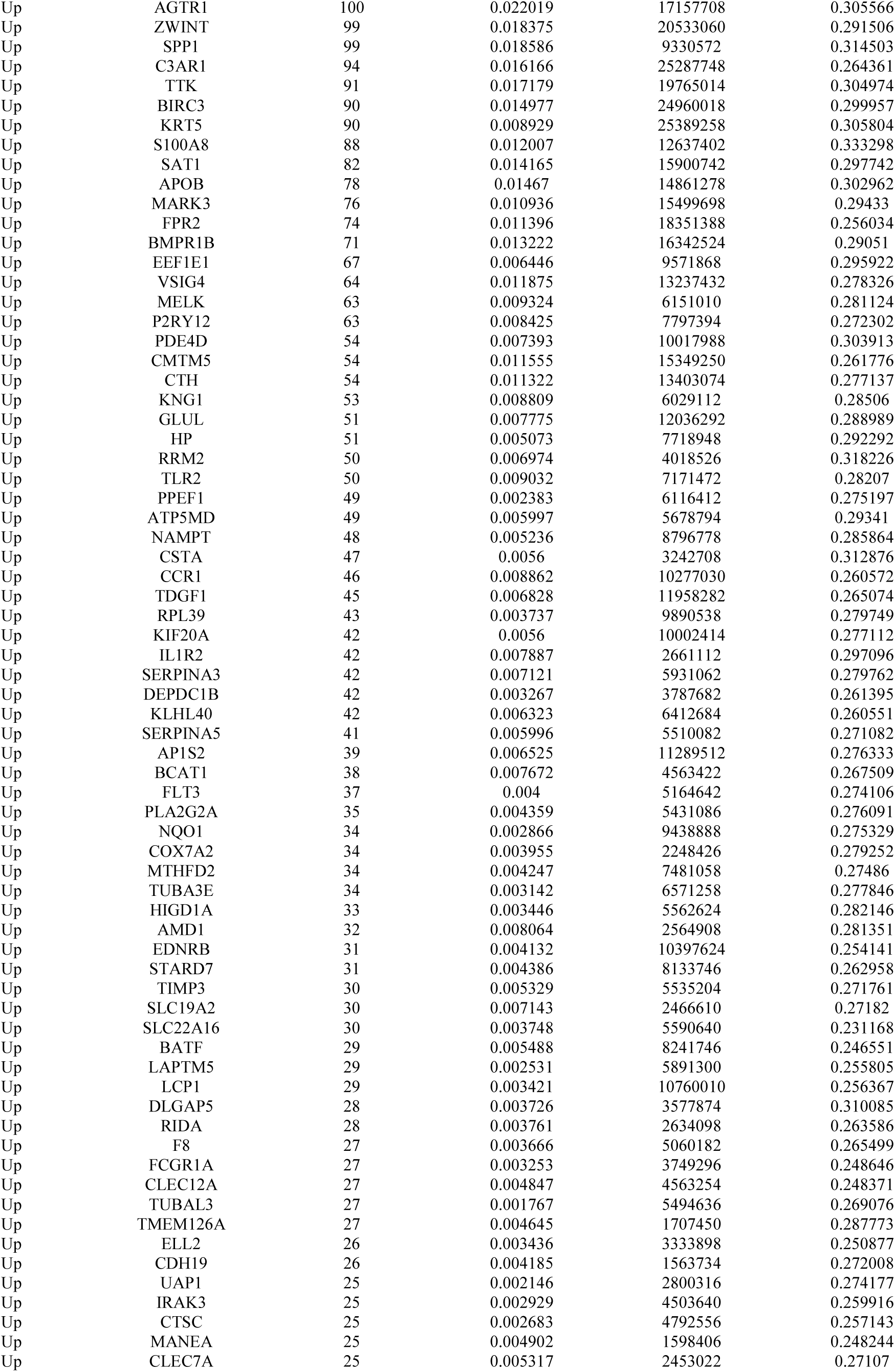
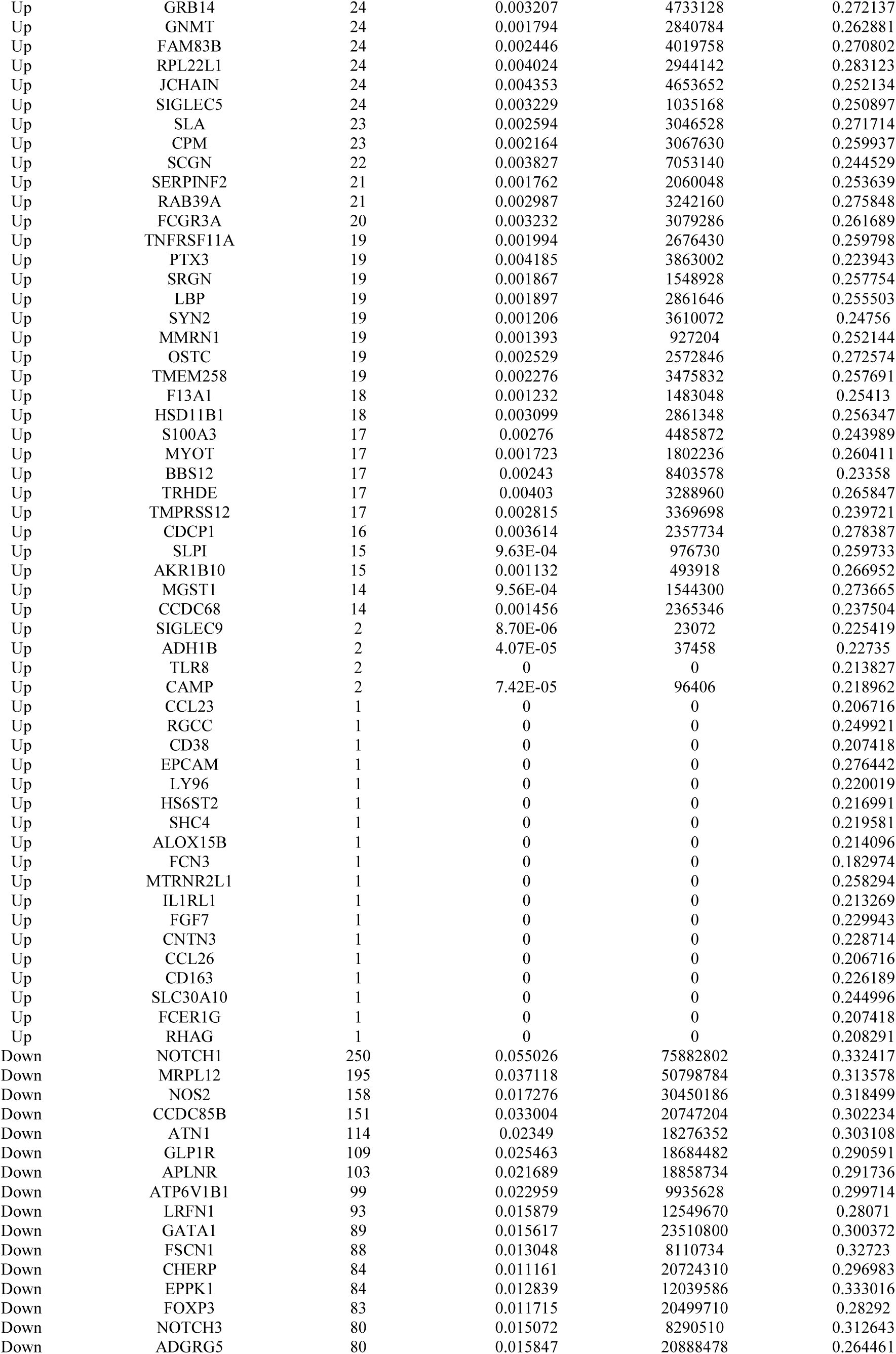
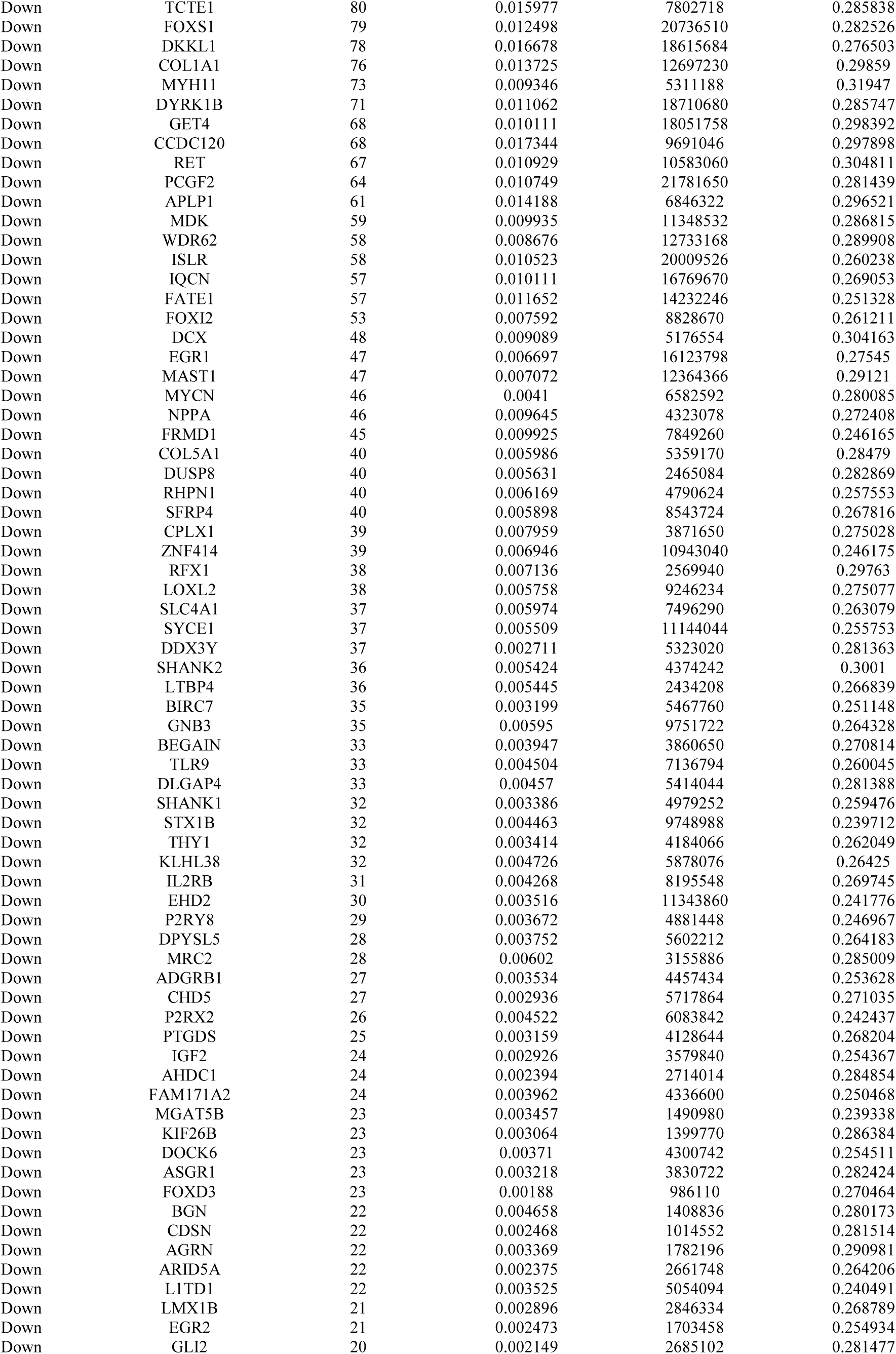
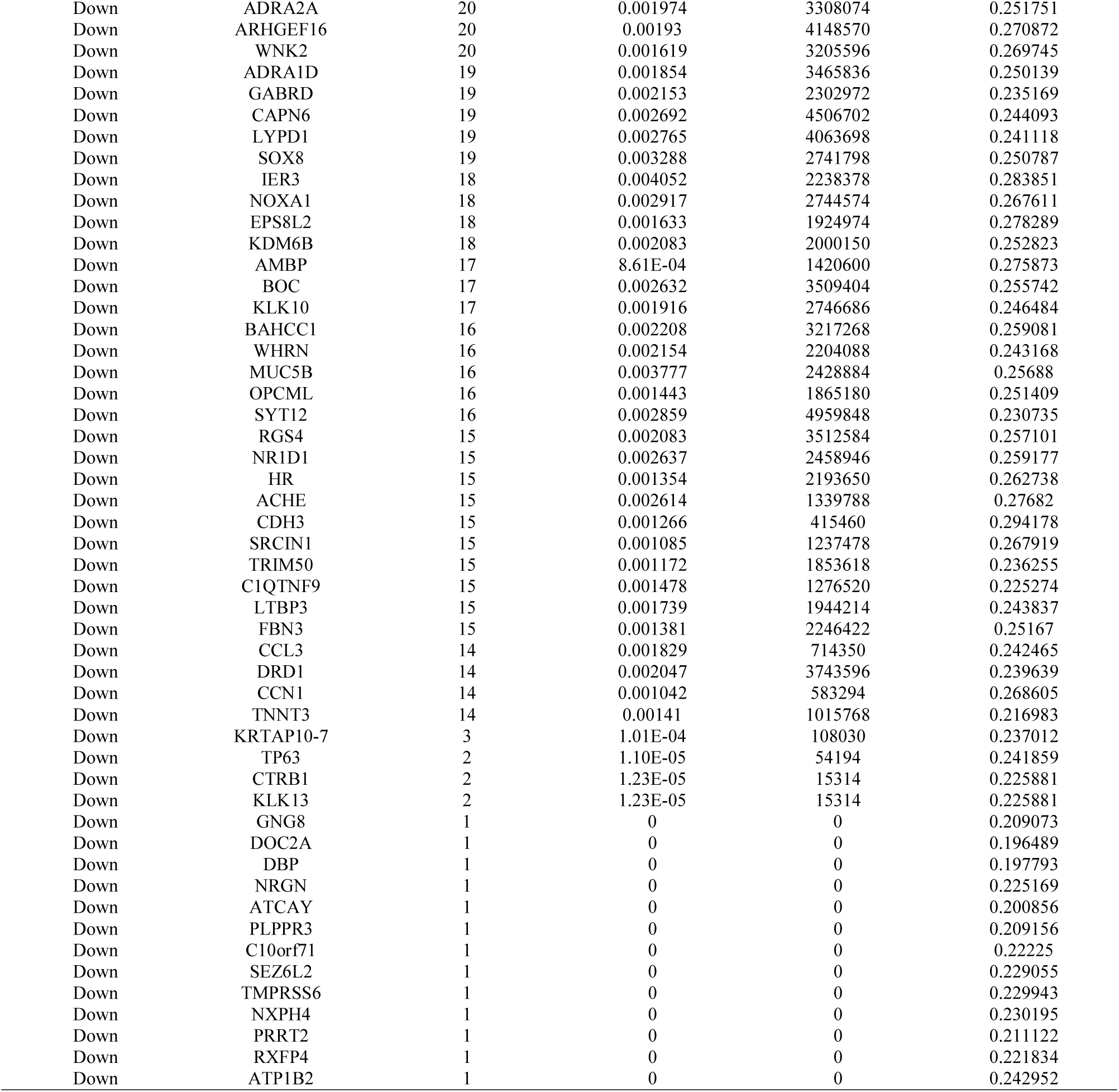
Topology table for up and down regulated genes.

### MiRNA-hub gene regulatory network construction

2222 nodes (miRNA: 1952 and hub gene: 270) and 9722 edges were mapped in the miRNA-hub gene regulatory network (Fig. 5). Moreover, we found that TFRC was potentially targeted by 108 miRNAs (ex; hsa-mir-409-3p); TOP2A was potentially targeted by 90 miRNAs (ex; hsa-mir-320d); CDK1 was potentially targeted by 68 miRNAs (ex; hsa-mir-301a-5p); CEP55 was potentially targeted by 56 miRNAs (ex; hsa-mir-107); FKBP5 was potentially targeted by 32 miRNAs (ex; hsa-mir-205-5p); ATN1 was potentially targeted by 74 miRNAs (ex; hsa-mir-3200-3p); FSCN1 was potentially targeted by 62 miRNAs (ex; hsa-mir-29a-5p); CHERP was potentially targeted by 52 miRNAs (ex; hsa-mir-296-5p); NOTCH1 was potentially targeted by 35 miRNAs (ex; hsa-mir-139-5p); NOTCH3 was potentially targeted by 33 miRNAs (ex; hsa-mir-147a) and are shown in Table 5.

**Fig. 5.**
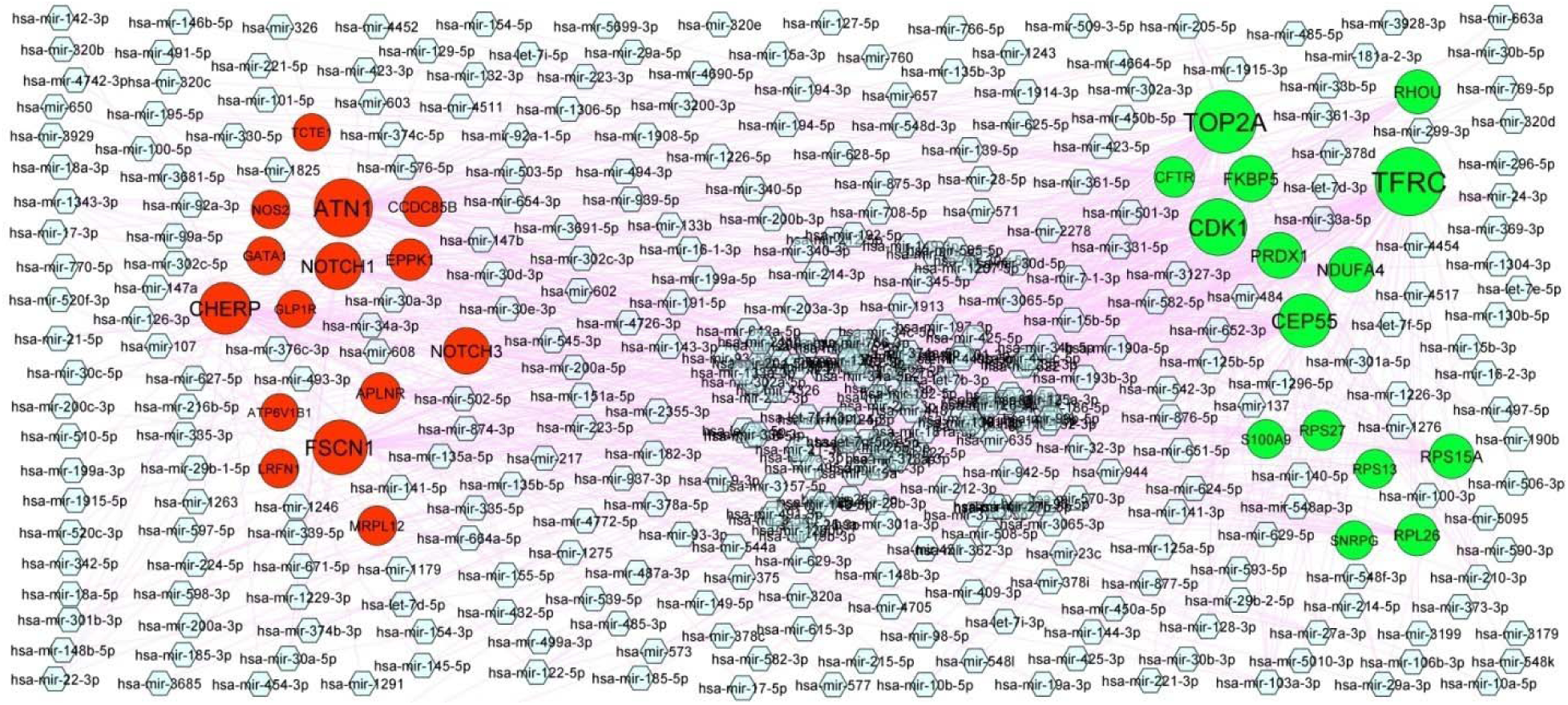
Target gene - miRNA regulatory network between target genes. The blue color diamond nodes represent the key miRNAs; up regulated genes are marked in green; down regulated genes are marked in red.

**Table 5.** miRNA - target gene and TF - target gene interaction

| Regulation | Target Genes | Degree | MicroRNA | Regulation | Target Genes | Degree | TF |
| --- | --- | --- | --- | --- | --- | --- | --- |
| Up | TFRC | 108 | hsa-mir-409-3p | Up | RPS27 | 18 | CREB1 |
| Up | TOP2A | 90 | hsa-mir-320d | Up | FKBP5 | 17 | TP63 |
| Up | CDK1 | 68 | hsa-mir-301a-5p | Up | TFRC | 15 | FOXA1 |
| Up | CEP55 | 56 | hsa-mir-107 | Up | CDK1 | 14 | NRF1 |
| Up | FKBP5 | 32 | hsa-mir-205-5p | Up | PRDX1 | 11 | JUND |
| Up | PRDX1 | 31 | hsa-mir-18a-3p | Up | CFTR | 10 | RUNX2 |
| Up | NDUFA4 | 28 | hsa-mir-185-5p | Up | RHOU | 10 | SOX17 |
| Up | RPS15A | 25 | hsa-mir-571 | Up | RPS13 | 10 | FOXF2 |
| Up | RHOU | 24 | hsa-mir-340-5p | Up | RPS15A | 9 | FOXC1 |
| Up | RPL26 | 17 | hsa-mir-200a-5p | Up | TOP2A | 8 | SRF |
| Up | CFTR | 12 | hsa-mir-4452 | Up | CEP55 | 7 | NFIC |
| Up | RPS27 | 11 | hsa-mir-103a-3p | Up | RPL26 | 7 | ZNF354C |
| Up | RPS13 | 7 | hsa-mir-126-3p | Up | SNRPG | 5 | ELK4 |
| Up | S100A9 | 7 | hsa-mir-1343-3p | Up | NDUFA4 | 3 | RELA |
| Up | SNRPG | 6 | hsa-mir-155-5p | Up | S100A9 | 2 | USF2 |
| Down | ATN1 | 74 | hsa-mir-3200-3p | Down | ATN1 | 19 | TEAD1 |
| Down | FSCN1 | 62 | hsa-mir-29a-5p | Down | FSCN1 | 8 | SREBF2 |
| Down | CHERP | 52 | hsa-mir-296-5p | Down | ATP6V1B1 | 7 | GATA2 |
| Down | NOTCH1 | 35 | hsa-mir-139-5p | Down | GATA1 | 7 | E2F1 |
| Down | NOTCH3 | 33 | hsa-mir-147a | Down | MRPL12 | 7 | TP53 |
| Down | EPPK1 | 16 | hsa-mir-148b-3p | Down | CHERP | 7 | PPARG |
| Down | APLNR | 13 | hsa-mir-373-3p | Down | GLP1R | 6 | ARID3A |
| Down | CCDC85B | 11 | hsa-mir-603 | Down | FOXP3 | 6 | STAT3 |
| Down | MRPL12 | 9 | hsa-mir-128-3p | Down | NOTCH3 | 6 | KLF5 |
| Down | LRFN1 | 6 | hsa-mir-145-5p | Down | EPPK1 | 5 | E2F6 |
| Down | NOS2 | 5 | hsa-mir-497-5p | Down | APLNR | 4 | PRDM1 |
| Down | GATA1 | 4 | hsa-mir-4454 | Down | NOS2 | 4 | IRF2 |
| Down | ATP6V1B1 | 2 | hsa-mir-1229-3p | Down | NOTCH1 | 3 | GATA3 |
| Down | GLP1R | 1 | hsa-mir-376c-3p | Down | LRFN1 | 3 | YY1 |
| Down | TCTE1 | 1 | hsa-mir-7-5p | Down | CCDC85B | 3 | TFAP2A |

### TF-hub gene regulatory network construction

356 nodes (TF: 78 and hub gene: 278) and 2152 edges were mapped in the TF-hub gene regulatory network (Fig. 6). Moreover, we found that RPS27 was potentially targeted by 18 TFs (ex; CREB1); FKBP5 was potentially targeted by 17 TFs (ex; TP63); TFRC was potentially targeted by 15 TFs (ex; FOXA1); CDK1 was potentially targeted by 14 TFs (ex; NRF1); PRDX1 was potentially targeted by 11 TFs (ex; JUND); ATN1 was potentially targeted by 19 TFs (ex; TEAD1); FSCN1 was potentially targeted by 8 TFs (ex; SREBF2); ATP6V1B1 was potentially targeted by 7 TFs (ex; GATA2); GATA1 was potentially targeted by 7 TFs (ex; E2F1); MRPL12 was potentially targeted by 7 TFs (ex; TP53) and are shown in Table 5.

**Fig. 6.**
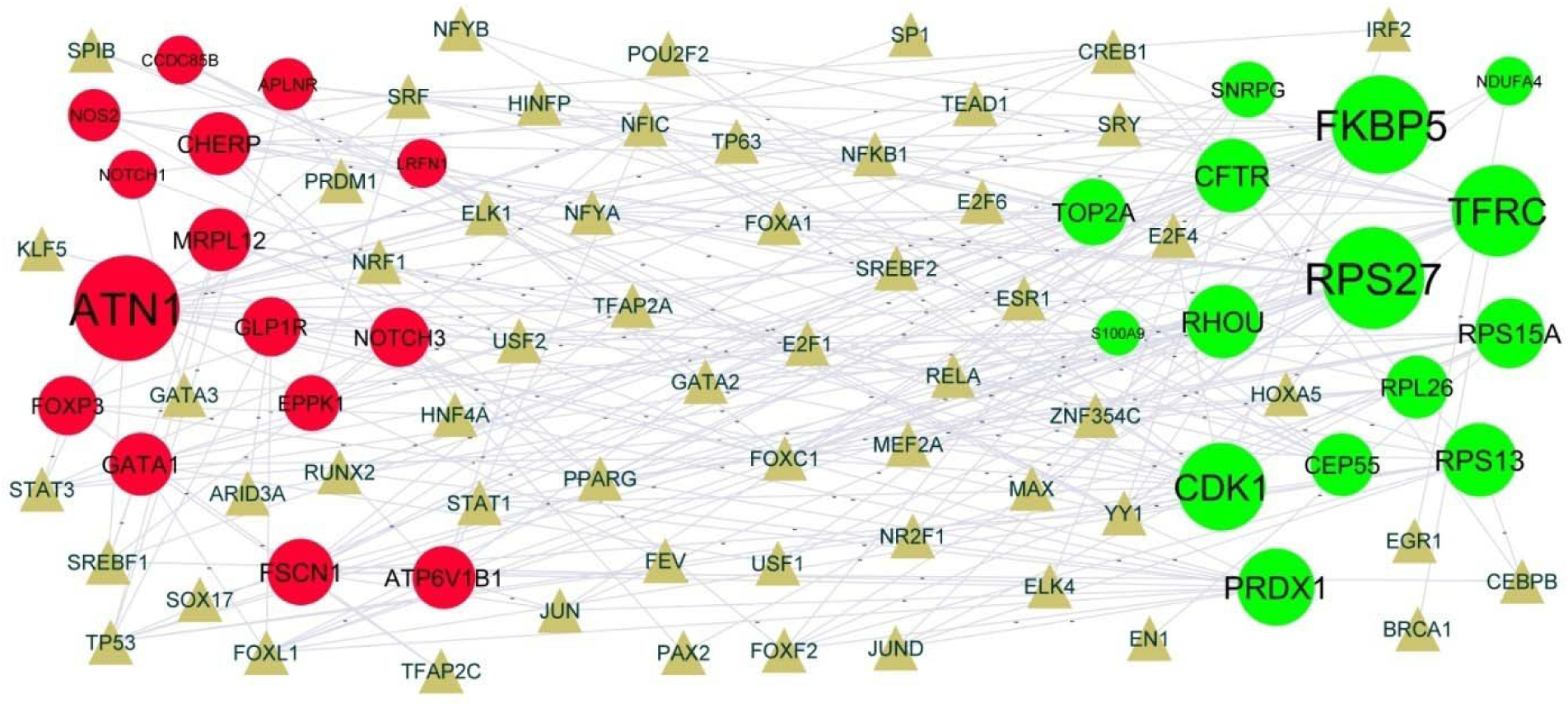
Target gene - TF regulatory network between target genes. The gray color triangle nodes represent the key TFs; up regulated genes are marked in green; down regulated genes are marked in red.

### Validation of hub genes by receiver operating characteristic curve (ROC) analysis

As these 10 hub genes are prominently expressed in MI, we performed a ROC curve analysis to evaluate their sensitivity and specificity for the diagnosis of MI. As shown in Fig. 7, CFTR, CDK1, RPS13, RPS15A, RPS27, NOTCH1, MRPL12, NOS2, CCDC85B and ATN1 achieved an AUC value of >0.88, demonstrating that these hub genes have high sensitivity and specificity for MI diagnosis. The results suggested that CFTR, CDK1, RPS13, RPS15A, RPS27, NOTCH1, MRPL12, NOS2, CCDC85B and ATN1 can be used as biomarkers for the diagnosis of MI.

**Fig. 7.**
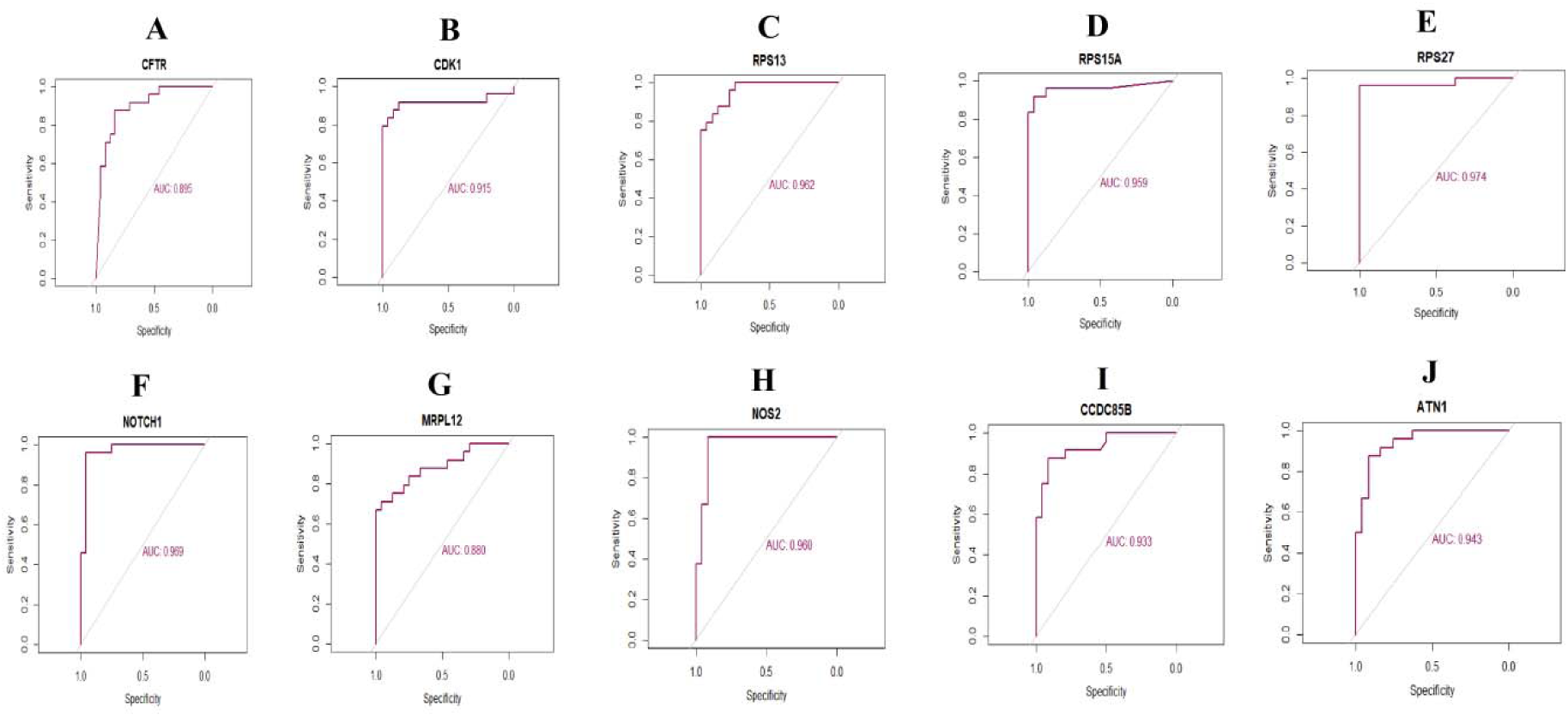
ROC curve analyses of hub genes. A) CFTR B) CDK1 C) RPS13 D) RPS15A E) RPS27 F) NOTCH1 G) MRPL12 H) NOS2 I) CCDC85B J) ATN1

## Discussion

Although people have continuously investigated MI, the early diagnosis and treatment of MI is still a large problem due to the lack of understanding of the molecular mechanisms that drive the occurrence and development of MI. Therefore, in-depth investigation into the factors and mechanisms of MI development are necessary for MI diagnosis and treatment. Due to well-developed RNA sequencing, it is easier to determine the general genetic modification in the progression of diseases, which can allow for the recognition of gene targets for diagnosis, therapy, and prognosis of MI.

In our investigation, a total of 958 DEGs were screened, including 480 up regulated genes and 478 down regulated genes. Lin et al [40] and Wuest et al [41] found that the role of IL1RL1 and ALOX15B in coronary artery disease. Research have shown that SERPINA3 [42], GPR78 [43] and ESM1 [44] plays a significant role in MI progression. Expression of the SCGN (secretagogin, EF-hand calcium binding protein) gene plays a role in the development of diabetes mellitus [45], but this genes might be novel target for MI.

In the present investigation, a GO and pathway enrichment analysis was conducted to further understand the regulatory roles of DEGs in MI. Immune system [46], innate immune system [47], neutrophil degranulation [48], toll-like receptor cascades [49], metabolism of amino acids and derivatives [50], neuronal system [51], extracellular matrix organization [52], degradation of the extracellular matrix [53], diseases of glycosylation [54], response to stimulus [55], cell communication [56], cell periphery [57], membrane [58], anatomical structure development [59] and plasma membrane [60] were responsible for progression of MI. PLA2G2A [61], CCL23 [62], CD53 [63], TREML4 [64], TREM2 [65], CD180 [66], HPSE (heparanase) [67], CELA2A [68], TNFRSF4 [69], AMBP (alpha-1-microglobulin/bikunin precursor) [70], SOX18 [71], PANX2 [72], RSPO2 [73], COMP (cartilage oligomeric matrix protein) [74], ASGR1 [75] and NOXA1 [76] are involved in progression of atherosclerosis. A previous study reported that S100A9 [77], ADORA3 [78], IL1R2 [79], FPR1 [80], CCL20 [81], CD163 [82], S100A8 [83], TLR2 [84], HAS2 [85], PTX3 [86], TIMP4 [87], AREG (amphiregulin) [88], LBP (lipopolysaccharide binding protein) [89], IL18R1 [90], ALOX5AP [91], RETN (resistin) [92], F13A1 [93], FPR2 [94], SAA1 [95], FLT3 [96], AQP4 [97], FCER1G [98], CCL18 [99], HP (haptoglobin) [100], CDK1 [101], SLC7A11 [102], CFTR (CF transmembrane conductance regulator) [103], F8 [104], STC1[44], IL18RAP [90], TIMP3 [105], PDE4D [106], CYP4A11 [107], SCN10A [108], APOB (apolipoprotein B) [109], ACE (angiotensin I converting enzyme) [110], PENK (proenkephalin) [111], HSPB6 [112], TLR9 [113], EGR1 [114], CACNG8 [115], FOXD3 [116], DBH (dopamine beta-hydroxylase) [117], FOXP3 [118], GLP1R [119], IL34 [120], CCN1 [121], ADRA2A [122], BGN (biglycan) [123], NOS2 [124], AGRN (agrin) [125], DRD1 [126], GNB3 [127], EGR2 [128], MDK (midkine) [129], NOTCH3 [130], AZIN2 [131], NOTCH1 [132], LOXL2 [133], ADAMTS14 [134] and SOD3 [135] are expressed in MI. DEFB1 [136], SLC11A1 [137], SPINK1 [138], CCR1 [139], GLUL (glutamate-ammonia ligase) [140], GPR84 [141], SIGLEC5 [142], SIGLEC7 [143], LGR5 [144], CD38 [145], GRB14 [146], PRDX1 [147], SLC19A2 [148], CADM2 [149], TRPM5 [150], COL1A1 [151], CTRB1 [152], UTS2R [153], CRTC1 [154], MUC5B [155], TMPRSS6 [156], SLC5A2 [157] and KCNJ9 [158] expression were shown to play an important role in diabetes mellitus, but these genes might be novel target for MI. MT1A [159], LYVE1 [160], S100A12 [161], GCKR (glucokinase regulator) [162], TLR8 [163], MRC1 [164], AGTR1 [165], P2RY12 [166], MSR1 [167], NQO1 [168], FKBP5 [169], CMTM5 [170], ADH1C [171], APLNR (apelin receptor) [172], SFRP4 [173], CCL3 [174], COL11A2 [175], EGR3 [176] and IL2RB [177] expression have a role in coronary artery disease. EDN2 [178], SNX10 [179] and KCNN1 [180] were played a predominant role in progression of atrial fibrillation. EDNRB (endothelin receptor type B) [181], FGF10 [182], WNK3 [183], KNG1 [184], FCN3 [185], AQP3 [186], HPR (haptoglobin-related protein) [187], CTH (cystathionine gamma-lyase) [188], SPARCL1 [189], MAOA (monoamine oxidase A) [190], BMPR1B (bone morphogenetic protein receptor type 1B) [191], FGF7 [192], CALCRL (calcitonin receptor like receptor) [193], MARK3 [194], ADH1B [195], AMH (anti-Mullerian hormone) [196], RET (ret proto-oncogene) [197], IGF2 [198], SLC6A9 [199], NPPA (natriuretic peptide A) [200], SCT (secretin) [201], DCX (doublecortin) [202], ASIC1 [203], LMX1B [204], DBP (D-box binding PAR bZIP transcription factor) [205] and SLC6A9 [206] levels are correlated with disease severity in patients with hypertension. CAMP (cathelicidin antimicrobial peptide) [207], SPP1 [208]. CXCL11 [209], TFRC (transferrin receptor) [210], IRAK3 [211], C3AR1 [212], GLI2 [213], THY1 [214], FOXO6 [215], RGS4 [216], WNT10B [217] and AEBP1 [218] were found to be involved in progression of obesity, but these genes might be novel target for MI. TDGF1 [219], KIF20A [220] and LTBP2 [221] were linked with progression of congenital heart defect.

A PPI network was constructed for the identified DEGs and hub genes were defined by the betweenness centrality, stress centrality and closeness centrality rank. The key modules were subsequently isolated from the PPI network. RPS13, RPS15A, RPS27, RPL26, RPS29, MRPL12, CCDC85B and ATN1 were the novel biomarkers for the progression of T2DM.

The subsequent construction of the miRNA-hub gene regulatory network and TF-hub gene regulatory network using the respective hub gens identified miRNA and TFs as potential key markers involved in MI. Chen et al [222], Li and Zhang [223], Wang et al [224], Zhao et al [225], Lin et al [226], Liao et al [227], Izadpanah et al [228], Wang et al [229] and Hakobjanyan et al [230] reported that the expression of the hsa-mir-409-3p, hsa-mir-320d, hsa-mir-107, hsa-mir-139-5p, NRF1, TEAD1, GATA2, E2F1 and TP53 are correlated with disease grades of MI. hsa-mir-301a-5p [231], hsa-mir-29a-5p [232], hsa-mir-296-5p [233], CREB1 [234] and FOXA1 [235] were linked with progression of diabetes mellitus, but these genes might be novel target for MI. hsa-mir-205-5p was involved in the progression of hypertension [236]. SREBF2 was associated with progression of coronary artery disease [237]. TOP2A, CEP55, FSCN1, CHERP, ATP6V1B1, GATA1, hsa-mir-3200-3p, hsa-mir-147a, TP63 and JUND (jun D proto-oncogene) were the novel biomarkers for the progression of MI.

In conclusion CFTR, CDK1, RPS13, RPS15A, RPS27, NOTCH1, MRPL12, NOS2, CCDC85B and ATN1 might be a critical gene for MI. In addition, immune system, neuronal system, response to stimulus, cell periphery, molecular transducer activity, multicellular organismal process and inorganic molecular entity transmembrane transporter activity play a pivotal role in the progression of MI from normal samples. Therefore, these pathways and genes can be a potential therapeutic target for target therapy in MI. We also introduce several miRNAs and TFs that can use to target these critical genes and pathways.

## Acknowledgement

I thank Juan Pablo Romero, CIMA, Hemato-Oncology, Advanced Genomics, Pamplona, Navarra, Spain, very much, the author who deposited their profiling by high throughput sequencing dataset GSE132143, into the public GEO database.

## Ethical approval

This article does not contain any studies with human participants or animals performed by any of the authors.

## Informed consent

No informed consent because this study does not contain human or animals participants.

## Availability of data and materials

The datasets supporting the conclusions of this article are available in the GEO (Gene Expression Omnibus) (https://www.ncbi.nlm.nih.gov/geo/) repository. [(GSE132143) https://www.ncbi.nlm.nih.gov/geo/query/acc.cgi?acc=GSE132143)]

## Consent for publication

Not applicable.

## Author Contributions

B. V. - Writing original draft, and review and editing

C. V. - Software and investigation

